# The immediate early protein 1 of human herpesvirus 6B counteracts ATM activation in an NBS1-dependent manner

**DOI:** 10.1101/2021.07.31.454588

**Authors:** Vanessa Collin, Élise Biquand, Vincent Tremblay, Élise G. Lavoie, Julien Dessapt, Andréanne Blondeau, Annie Gravel, Louis Flamand, Amélie Fradet-Turcotte

**Author notes:** Co-corresponding authors **Co-corresponding authors’ information:** Amélie Fradet-Turcotte, CRCHU de Québec Research center – Hotel-Dieu de Québec, 9 McMahon, room 3744, Québec, Québec, Canada, G1R 2J6, Louis Flamand, CRCHU de Québec Research center – CHUL, 2705 boulevard Laurier, T1-64, Québec, Québec, Canada G1V 4G2. These authors contributed equally to this work (Vanessa Collin, Élise Biquand, Vincent Tremblay). **Author Contributions:** V.C., E.B., V.T., L.F., and A.F.-T. designed research; V.C., E.B., V.T., E.G.L., J.D., A.B., and A.G. performed research and analyzed data; V.C. and A.F.-T. wrote the original draft and E.B., V.T., and L.F. edited the manuscript.

## Abstract

Viral infection often trigger an ATM-dependent DNA damage response (DDR) in host cells that suppresses viral replication. To counteract this antiviral surveillance system, viruses evolved different strategies to induce the degradation of the MRE11/RAD50/NBS1 (MRN) complex and prevent subsequent DDR signaling. Here, we report that human herpesvirus 6B (HHV-6B) infection causes genomic instability by suppressing the host cell’s ability to induce ATM-dependent signaling pathways. Expression of immediate early protein 1 (IE1) phenocopies this phenotype and blocks further homology-directed double-strand break (DSB) repair. In contrast to other viruses, IE1 does not affect the stability of the MRN complex. Instead, it uses two distinct domains to inhibit ATM serine/threonine kinase (ATM) activation at DSBs. Structure-based analyses revealed that the N-terminal domain of IE1 interacts with the BRCA1 C-terminal domain 2 of nibrin (NBN, also known as NBS1), while ATM inhibition is attributable to on its C-terminal domain. Consistent with the role of the MRN complex in antiviral responses, NBS1 depletion resulted in increased HHV-6B replication in infected cells. However, in semi-permissive cells, viral integration of HHV-6B into the telomeres was not strictly dependent on NBS1, supporting models where this process occurs via telomere elongation rather than through DNA repair. Interestingly, as IE1 expression has been detected in cells of subjects with inherited chromosomally-integrated form of HHV-6B (iciHHV-6B), a condition associated with several health conditions, our results raise the possibility of a link between genomic instability and the development of iciHHV-6-associated diseases.

**Significance Statement:** Many viruses have evolved ways to inhibit DNA damage signaling, presumably to prevent infected cells from activating an antiviral response. Here, we show that this is also true for human herpesvirus 6B (HHV-6B), through its immediate early protein 1 (IE1). However, in contrast to adenovirus’ immediate early proteins, HHV-6B IE1 is recruited to double-strand breaks in an NBS1-dependent manner and inhibits ATM serine/threonine kinase activation. Characterizing this phenotype revealed a unique mechanism by which HHV-6B manipulates DNA damage signaling in infected cells. Consistently, viral replication is restricted by the MRN complex in HHV-6B infected cells. Viral integration of HHV-6B into the host’s telomeres is not strictly dependent on NBS1, challenging current models where integration occurs through homology-directed repair.

## Introduction

To infect a cell, a virus needs to successfully replicate its genetic material and produce new virions. Accordingly, cells have a sophisticated surveillance system that detects viral DNA and activates an innate antiviral response. Mounting evidence supports a role for the DNA damage response (DDR) in this process, revealing an intricate interplay between its activation and the activation of intrinsic antiviral responses (1). Some viruses, such as adenovirus, target the MRE11-RAD50-NBS1 (MRN) complex for degradation to prevent the activation of ATM serine/threonine kinase (ATM) (2, 3), while others, such as herpes simplex virus 1 (HSV-1) and human papillomavirus, rely on these proteins for efficient viral replication (4–6). How and why viruses inhibit or hijack the ATM pathway remains a mystery (7).

In mammalian cells, the MRN complex and ATM are essential to maintain genomic stability in the presence of DNA double-strand breaks (DSBs). Broken DNA ends are first detected by the MRN complex (8), where its accumulation induces a signaling cascade that activates ATM and subsequent phosphorylation of the histone variant H2AX on Ser139 (producing γ-H2AX). Mediator of DNA damage checkpoint 1 (MDC1) interacts with γ-H2AX, triggering the ubiquitylation of chromatin surrounding the break by promoting the accumulation of the E3-ubiquitin ligases ring finger protein (RNF) 8 and RNF168 (9, 10). In the G1 phase of the cell cycle, the recruitment of the DNA repair factor tumor protein p53 binding protein 1 (53BP1) at ubiquitylated chromatin promotes DNA repair *via* non-homologous end-joining (NHEJ)(11). In the S and G2 phases, BRCA1 DNA repair associated (BRCA1) and RB binding protein 8 endonuclease (CtIP) accumulate at the break and cooperate with exonuclease 1 (EXO1), BLM RecQ like helicase, and DNA replication helicase/nuclease 2 (BLM-DNA2) to facilitate end resection. This process results in extensive single-stranded (ss) DNA accumulation, which ultimately triggers the recruitment of recombinases that drive the homology searching required for homology-driven recombination (HDR) (12). HDR uses homologous sequences as templates to repair breaks in a faithful manner and includes processes such as homologous recombination (HR), single-stranded annealing (SSA), and break-induced replication (BIR) (11, 13–15).

Human herpesvirus 6B (HHV-6B) is a betaherpesvirus that infects nearly 90% of the world’s population in the first 2 years of life and is responsible for roseola infantum, a pathology defined by skin rashes, high fever, and respiratory distress (16–18). In this double-stranded DNA (dsDNA) virus subfamily, HHV-6B shares 90% homology with HHV-6A, another lymphotropic virus. Although both infect CD4^+^ T lymphocytes, they have epidemiological, biological, and immunological differences (19). Like other herpesviruses, HHV-6A and HHV-6B (HHV-6A/B) establish lifelong latency in infected hosts and can occasionally reactivate (20). However, whereas most herpesviruses achieve latency by circularizing and silencing their genome, HHV-6A/B integrate their genomes into the host’s chromosomal terminal repeats (telomeres) (21, 22). The linear dsDNA genomes of HHV-6A/B are both flanked by an array of direct repeats containing 15–180 reiterations of perfect telomeric repeats (pTMRs) identical to the human telomeric sequence (5′-TTAGGG-3′), enabling viral integration (23). This process depends on the integrity of these pTMRs (23), resulting in a model in which viral integration is mediated through HDR, including the SSA or BIR DNA repair mechanisms (24). When HHV-6A/B integration occurs in a gamete before fertilization, the newborn carries a copy of HHV-6A/B in every cell of its body and can be transmitted to its offspring. This condition, called inherited chromosomally-integrated (ici)HHV-6A/B, affects ∼1% of the world’s population, representing almost 80 million people (25, 26). It is more prevalent in those suffering from health issues such as high spontaneous abortion rates (27, 28), pre-eclampsia (29), and angina pectoris (30) compared to healthy subjects (reviewed in (31, 32)). However, the reason why iciHHV-6A/B contributes to these clinical syndromes has not been elucidated in any detail.

HHV-6B, which is better characterized than HHV-6A, sequentially expresses more than 97 genes/proteins during its lytic cycle(33). Immediate early (IE) proteins are expressed early in the viral cycle, to regulate viral gene expression and establish a favorable environment for viral replication. In the context of HHV-6B, IE protein 1 (IE1) is the first protein expressed during cell infection (34). It inhibits the innate antiviral response in part by sequestering signal transducer and activator of transcription 2 (STAT2) in the nucleus, thereby compromising type I interferon production and signaling(35, 36). In infected cells, IE1 is exclusively localized within PML bodies (37), which were recently implicated in HDR-mediated DNA repair through an undefined mechanism (38–41). Interestingly, PML depletion reduces HHV-6B integration (42), suggesting that IE1-containing PML bodies also participate in viral integration.

In this study, we report that HHV-6B infection—and more specifically, IE1 expression—leads to genomic instability in cells. Further investigations revealed that IE1 specifically prevents phosphorylation of the histone variant H2AX and subsequent HDR repair. Structure-function analyses reveal that IE1 interacts with NBS1 and inhibits ATM. Consistent with a role for the MRN complex in interfering with viral replication, we show that NBS1 depletion results in increased HHV-6B replication in infected cells. Although current models propose that viral integration occurs through HDR DNA repair, we show that viral integration is not affected in NBS1-depleted cells that elongate their telomeres in a human telomerase reverse transcriptase (hTERT)-dependent manner (42). Thus, our findings reveal that viral integration relies on biological pathways that safeguard telomere extension in infected cells and not on specific DNA repair pathways.

## Results

### HHV-6B infection and IE1 expression induce genomic instability

Our first indication that HHV-6B infection induces genomic instability in host cells was the observation that HHV-6B infection rapidly induced micronuclei (MNi) formation in MOLT-3 cells (a lymphoblast T cell line; Fig.1*A* and *SI Appendix*, Fig. S1*A*). In these experiments, cells infected with the HHV-6B strain Z29 accumulated 6.6-fold more MNi than non-infected cells (Mock) 24 h post-infection (Fig.1*A*). Consistent with the rapid accumulation of genomic instability in infected cells, we observed a similar phenotype in duplicate clones of stable U2OS cell lines containing a doxycycline (Dox)-inducible expression cassette for IE1 (C10 and C102; Fig. 1*B* and *SI Appendix*, Fig. S1*B-C*). These clones had 4.8-fold more MNi than parental U2OS cells 48 h post-IE1 induction (Fig.1*B*), suggesting that the instability observed in HHV-6B infected cells is at least partially caused by IE1. Note that both U2OS clone (C10 and C102) exhibit similar levels of MNi than the parental cell line without IE1 induction (*SI Appendix*, Fig. S1*C*). MNi arise from unresolved genomic instabilities such as DSBs (i), lagging chromosomes (ii), and ruptured anaphase bridges (ABs) (iii) (Fig. 1*C*) (43). They are compartmentally separated from the primary nucleus and surrounded by a nuclear envelope, as shown by the presence of lamin B in these perinuclear structures (*SI Appendix*, Fig. S1*D-E*) (44). To determine how IE1 triggers MNi formation, we analyzed the accumulation of different markers in IE1-induced MNi, such as centromeres and telomeric DNA (to detect lagging chromosomes and ABs caused by telomere fusion, respectively). Interestingly, a much lower proportion of the IE1-induced MNi contained centromeres compared with those in parental U2OS cells (∼10-fold, Fig. 1*D* and *SI Appendix*, Fig. S1*F*), suggesting that they are not induced by chromosome segregation defects. Although IE1 partially colocalizes with telomeres in host cells (42), fluorescence *in situ* hybridization (FISH) revealed that IE1 does not specifically promote instability at telomeres, as IE1-induced MNi accumulated similar levels of telomeric DNA as those in parental U2OS cells (Fig. 1*E* and *SI Appendix*, Fig. S1*G*). Metaphase spread assays revealed that IE1-expressing U2OS cell lines exhibited higher frequencies of DNA breaks than parental cells (Fig. 1*F-G*), consistent with MNi being induced by DSB accumulation. Interestingly, IE1 was detected in only 5–10% of the MNi (*SI Appendix*, Fig. S1*H*), suggesting that they are not arising from DSBs induced by the physical binding of IE1 at any defined DNA locus.

**Fig. 1.**
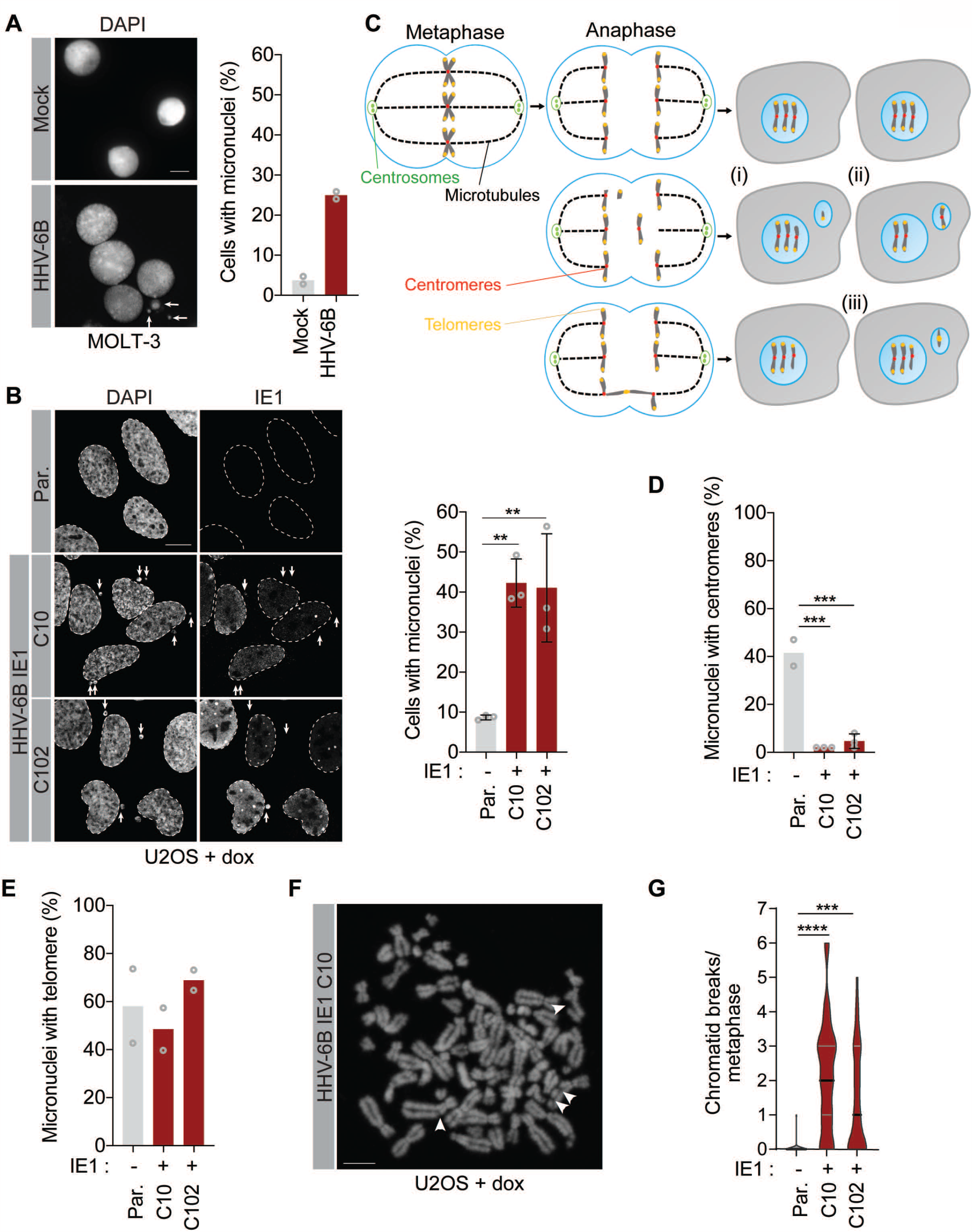
HHV-6B infection and IE1 expression lead to micronuclei formation *(A)* Left panel: Representative images of Mock and HHV-6B-infected MOLT-3 cells fixed 24 h post-infection and counterstained with DAPI. Micronuclei are indicated by white arrows. The quantification of the micronuclei (right panel) shows the mean (*n* = 2, >100 micronuclei/condition). *(B)* Left panel: representative images of U2OS cell lines and clones stably expressing Dox-inducible HHV-6B IE1 (C10 and C102). IE1 expression was induced with 1 μg/mL Dox for 48 h prior to IE1 immunofluorescence. Micronuclei are indicated by white arrows. The parental cell line (Par.) was used as a negative control. The micronuclei (right panel) quantification shows the mean ± standard deviation (SD; *n* = 3). Schematic of micronuclei formation *via* DNA double-strand breaks (DSBs) (i), lagging chromosomes (ii), and anaphase bridges (ABs) (iii). *(D-E)* Quantification of micronuclei containing centromeres (D) and telomeres (E). Cells were treated as in B and centromeres and telomeres were detected by immunofluorescence and FISH, respectively. Data represent the mean ± SD (*n* = 3) (D) and the mean (*n* = 2, > 100 micronuclei/condition) (E). *(F)* A representative metaphase spread from an IE1-expressing cell. Cells were treated with 1 μg/mL Dox for 48 h, then metaphase spreads were prepared, fixed, and counterstained with DAPI. *(G)* Quantification of chromosomal aberrations per metaphase. Data represent the mean ± SD (*n* = 31). \*\**p* < 0.01, \*\*\**p* < 0.001, \*\*\*\**p* < 0.0001. Scale bars = 5 μm.

### HHV-6B IE1 impairs DSB signaling and homology-directed DNA repair

DSB accumulation results from either an increase in DNA breaks or defective DNA DSB signaling and repair. To determine how HHV-6B infection promotes genomic instability, we first investigated whether infected cells accumulated Ser139-phosphorylated H2AX (*i*.*e*., the DSB marker γ-H2AX). Surprisingly, while γ-H2AX foci accumulated in > 75% of non-infected MOLT-3 cells exposed to irradiation (IR), this number was dramatically reduced in infected cells (Fig. 2*A-B* and *SI Appendix*, Fig. S2*A*). U2OS clones stably expressing IE1 reproduced this phenotype (Fig. 2*C-D* and *SI Appendix*, Fig. S2*B*), indicating that IE1 impairs DSB signaling. This inhibition is independent of IE1 accumulation within PML bodies, as no γ-H2AX foci were detected in PML-deficient U2OS cells that transiently express IE1 (*PML*^−/−^, Fig. 2*E* and *SI Appendix*, Fig. S2*C-E*).

**Fig. 2.**
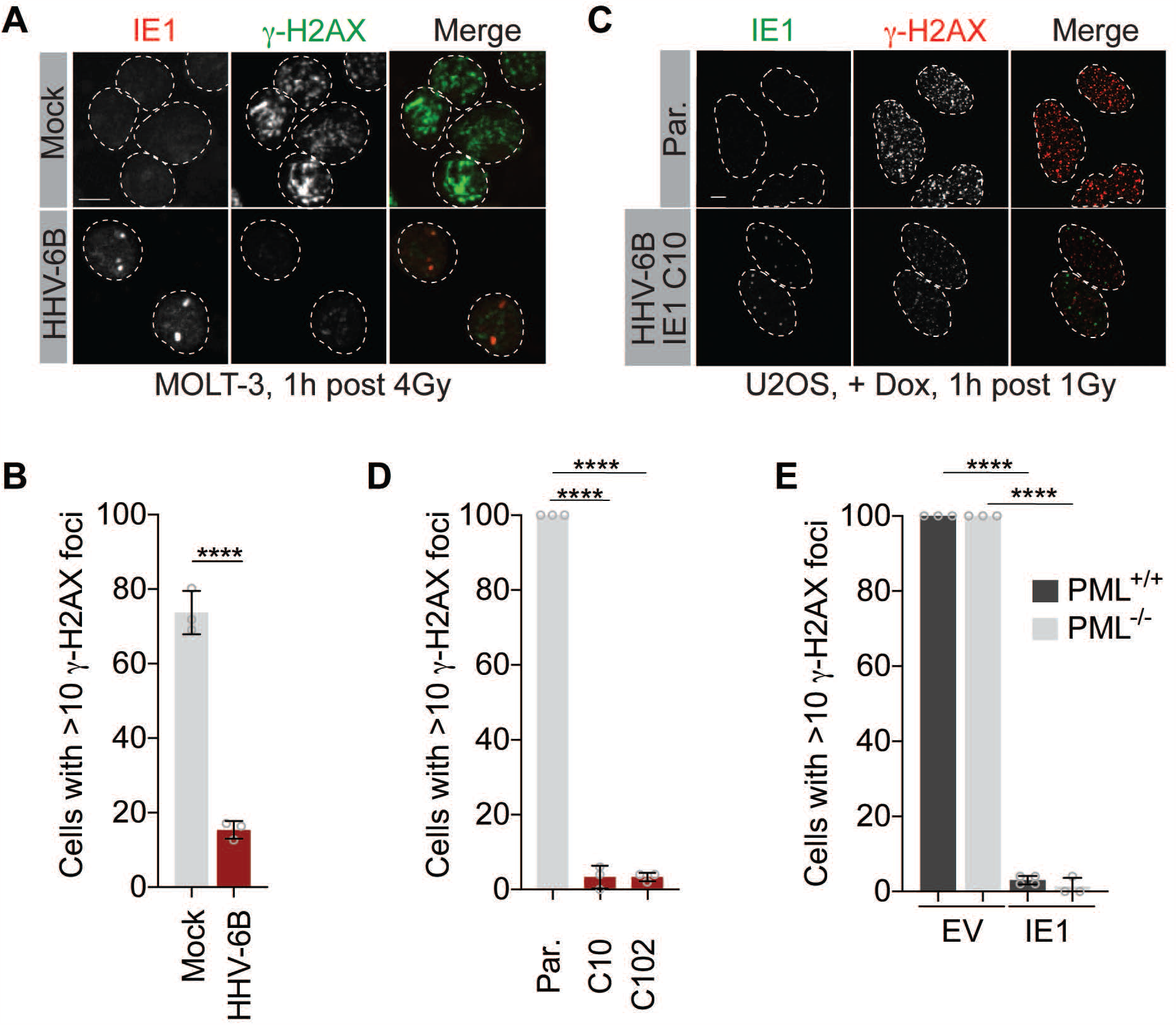
H2AX phosphorylation (γ-H2AX) is inhibited in HHV-6B-infected and IE1-expressing cells *(A)* Representative γ-H2AX immunostaining in HHV-6B-infected MOLT-3 cells irradiated with 4 Gy and immunostained for IE1 and γ-H2AX 1 h later. Mock-infected cells were used as a negative control. *(B)* Quantification of irradiated MOLT-3-infected and Mock cells with > 10 γ-H2AX foci. Data represent the mean ± SD (*n* = 3). Statistical significance was assessed using unpaired *t*-test. *(C)* Representative γ-H2AX immunostaining in irradiated U2OS parental (Par.) and IE1-expressing cells (Clone C10). IE1 expression was induced as in Fig. 1B. Cells were irradiated with 1 Gy and immunostained for IE1 and γ-H2AX 1 h later. *(D)* Quantification of irradiated U2OS Par. and IE1-expressing cells with > 10 γ-H2AX foci. Data are presented as the mean ± SD (*n* = 3). *(E)* Quantification of cells with > 10 γ-H2AX foci in irradiated (1 Gy) U2OS *PML*^+/+^ and ^−/-^ cells that transiently express untagged IE1. An empty vector (EV) was used as a negative control. Data represent the mean ± SD (*n* = 3). \*\*\*\**p* < 0.0001. Scale bars = 5 μm.

DSB signaling is essential to activate DNA repair pathways. Therefore, we tested whether DNA repair is inhibited in HHV-6B IE1-expressing cells. As mammalian cells use several pathways to repair DSBs (45), we performed a panel of DNA repair reporter assays in cells with stable or transient IE1 expression. These reporter assays all rely on the detection of a fluorescent protein that is expressed only if a site-specific DSB (induced by either I-SceI or Cas-9) is adequately repaired (Fig. 3*A-D* and *SI Appendix*, Fig. S3*A*, top panels) (46, 47). In the DR-GFP (direct repeats) and CRISPR-LMNA HDR assays, DSBs repaired by HR either reconstitute a defective green fluorescent protein (GFP)-reporter transgene integrated into the genome (DR-GFP) or introduce a cassette expressing mRuby in frame with endogenous lamin A (CRISPR-LMNA HDR), respectively (48, 49). In SA-GFP (single-strand annealing), proper annealing of a small homologous stretch reconstitutes a truncated GFP (48). In BIR-GFP (break-induced replication), replication-mediated repair following homology searching places a GFP coding sequence in the correct orientation (15, 50). Finally, in NHEJ-GFP, the ends of two DSBs need to be correctly ligated to recreate a full-length (FL) GFP-expressing cassette (51, 52). In assays using reporter transgenes integrated into the genome (DR-GFP, SA-GFP, BIR-GFP, and NHEJ-GFP) (Fig. 3*A-D* and *SI Appendix*, Fig. S3*B-C*), a condition without endonuclease I-SceI or Cas-9 was used as a negative control and the percentage of fluorescent cells obtained with I-SceI was set to 1. In each condition, a small amount of a near-infrared fluorescent protein (iRFP)-expressing vector was transfected with I-SceI, IE1, or an empty vector (EV) to ensure that DNA repair was measured in transfected cells only, and DNA repair was assessed 48 or 72 h post-transfection. Finally, as the clonal BIR-GFP U2OS cell line was generated in this study using a previously described BIR-GFP reporter plasmid(50), we used short interfering (si)RNAs against RAD51 and RAD52 as additional controls (*SI Appendix*, Fig. S3*D-F*) (13). As expected, the BIR-GFP signal was specifically inhibited in cells depleted of RAD51 (50).

**Fig. 3.**
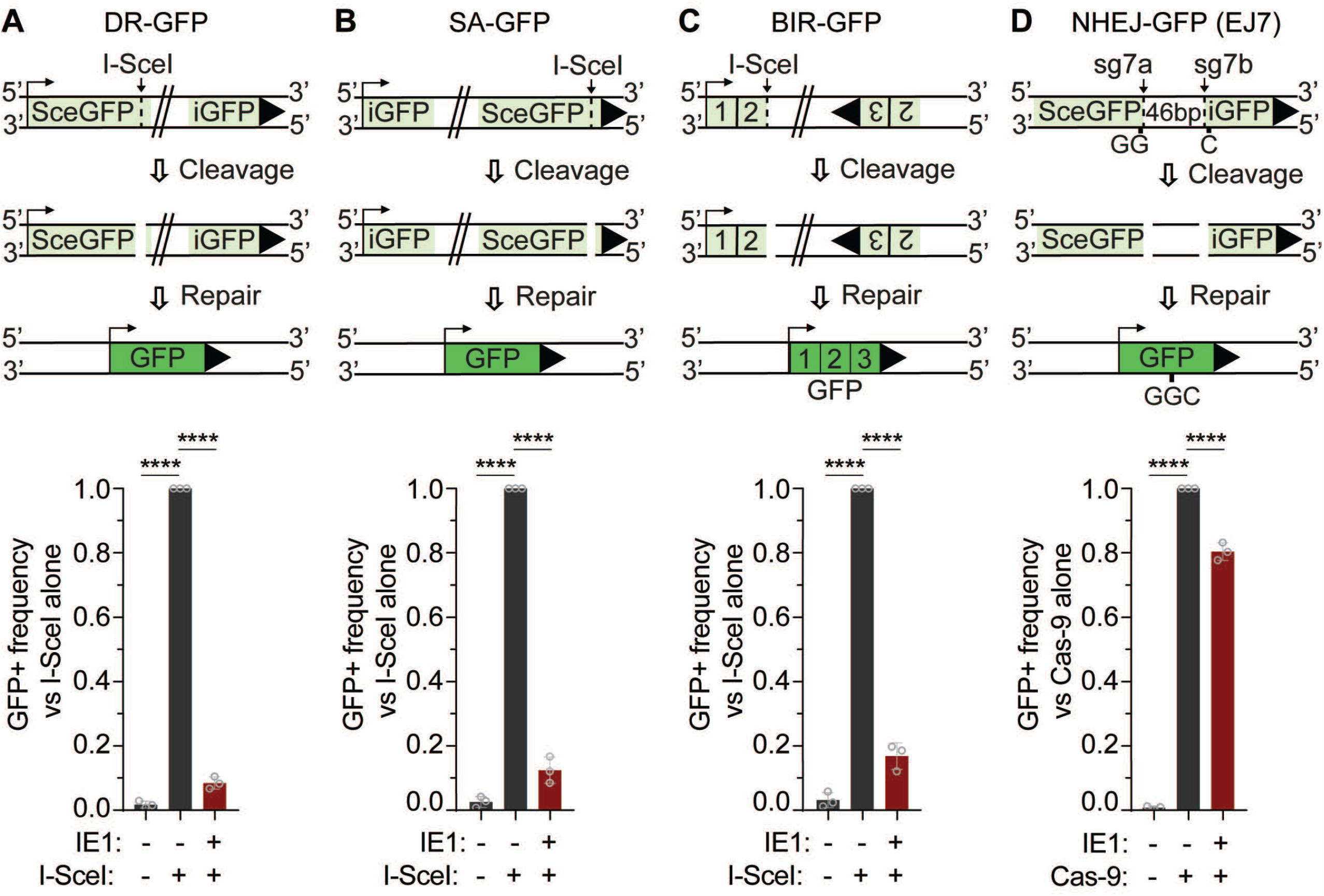
HHV-6B IE1 inhibits HDR-mediated repair *(A-D)* DNA repair reporter assays for (A) homologous recombination (DR-GFP), (B) single-strand annealing (SA-GFP), (C) break-induced replication (BIR-GFP), and non-homologous end-joining (NHEJ-GFP (EJ7)). For each assay, a schematic is presented in the top panel and the flow cytometry-based quantification of GFP^+^ cells is presented in the bottom panel. In each replicate, GFP^+^ cells were normalized to the GFP^+^ cells in the positive control (I-SceI^+^ or Cas9^+^, set to 1.0). Data represent the mean ± SD (*n* = 3). ****p < 0.0001.

Interestingly, these analyses revealed that both transient and stable IE1 expression drastically reduced all types of homology-directed DNA repair (Fig. 3*A-C*, lower panels, and *SI Appendix*, Fig. S3*A-C*). In contrast, IE1 only slightly modulated DNA repair in reporter assays assessing NHEJ (Fig. 3*D*). As the choice between HDR and NHEJ is driven by the cell cycle (45), we confirmed that cell cycle progression was not affected in cells expressing IE1 (*SI Appendix*, Fig. S3*G*). Altogether, these results show that homology-based DNA repair is specifically inhibited in cells expressing HHV-6B IE1.

### HHV-6B IE1 interacts with NBS1 and inhibits its ability to promote ATM activation

At DSBs, homology-based DNA repair is initiated when lesions are detected by the MRN complex, which leads to ATM auto-activation (Fig. 4*A*) through a still poorly understood mechanism (8, 53, 54). In adenovirus-infected cells, ATM activation is inhibited through the degradation of the MRN complex, which is mediated by the protein E4 (2). In contrast, MRN complex components were stable at steady state upon IE1 induction in U2OS clones (Fig. 4*B*). Interestingly, immunofluorescence analyses revealed that IE1 colocalizes with NBS1 in ∼75% of IE1-expressing cells (Fig. 4*C-E* and *SI Appendix*, Fig. S4*A-B*). This colocalization was also detected in irradiated cells and *PML* knockout cells (*SI Appendix*, Fig. S4*C-E*), suggesting that the interaction between IE1 and NBS1 is constitutive and independent of PML bodies. We also observed constitutive colocalization between MRE11 and IE1 (Fig. 4*D-E* and *SI Appendix*, Fig. S4*A-B, F*). However, the colocalization of transiently expressed IE1 with MRE11 was greatly reduced upon NBS1 depletion (Fig. 4*F* *and SI Appendix*, Fig. S4*G-H*), supporting a model where IE1 colocalizes with the MRN complex by interacting with NBS1. Importantly, we confirmed that colocalization between IE1 and NBS1 is also observed in HHV-6B-infected MOLT-3 cells (Fig. 4*G-H*).

**Fig. 4.**
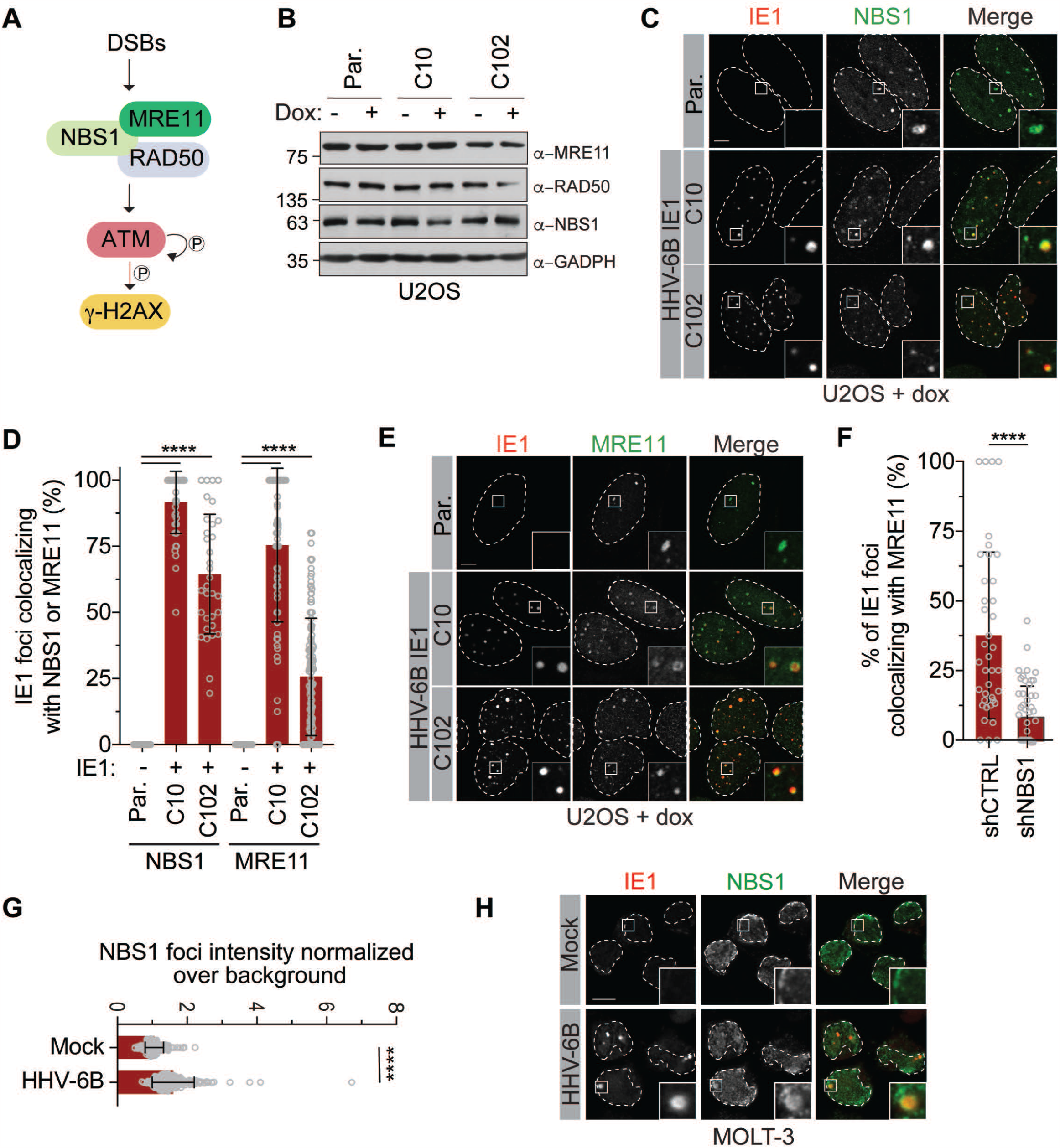
HHV-6B IE1 colocalizes with NBS1 *(A)* Signaling events triggered by DNA DSBs. *(B)* Whole cell extracts (WCEs) from U2OS cells (Par.) and IE1-expressing U2OS stable cell lines treated with or without 1 μg/mL Dox were immunoblotted for RAD50, NBS1, and MRE11. GAPDH was used as a loading control. *(C, E)* Representative images of the colocalization between IE1 and NBS1 (C) and MRE11 (E). IE1-expressing cells were treated as described in Fig. 1B and immunostained for IE1, NBS1, or MRE11. As a positive control, irradiated U2OS cells (+IR) were fixed 15 min post-irradiation (1 Gy) and immunostained as indicated (*SI Appendix*, Fig. S4*A*; scale bars, 5 μm). The parental cell line (Par.) was used as a negative control. *(D)* Percentages of IE1 foci that colocalized with NBS1 (C) and MRE11 (E). Data represent the mean ± SD of three independent experiments. *(F)* Percentage of IE1 foci that colocalized with MRE11 in stable U2OS control cells (shCTRL) or those depleted of NBS1 (shNBS1). Data represent the mean ± SD (*n* = 2, at least 40 nuclei/condition). Statistical significance was assessed using unpaired *t*-test. \*\*\*\**p* < 0.0001. Scale bars = 5 μm.

To further characterize the interplay between IE1 and the MRN complex, we took advantage of a cell-based assay that quantifies the ability of a mCherry-LacRnls fusion protein to specifically induce the recruitment of a “prey” to a *LacO* array integrated at a single locus in U2OS 2-6-5 cells (*(i)* No DSBs, Fig. 5A) (55, 56). This system can also be used to study signaling at DSBs by recruiting the ER-mCherry-LacR-FOKI-DD endonuclease to the *LacO* array (*(ii)* Localized DSBs, Fig. 5A). Although ER-mCherry-LacR-FOKI-DD is constitutively expressed in U2OS 2-6-5 cells, the protein is cytoplasmic, and a C-terminal destabilization domain (DD) ensures its continual degradation (55). DSBs can be rapidly induced by adding 4-hydroxytamoxifen (4-OHT) and Shield-1 to the culture medium. 4-OHT induces nuclear relocalization of ER-mCherry-LacR-FOKI-DD *via* its modified estrogen receptor (ER) domain and Shield-1 stabilizes it by inactivating the DD. When the mCherry-LacRnls-IE1 fusion protein was transiently expressed in U2OS 2-6-5 cells, only NBS1 was efficiently recruited to the *LacO* (Fig. 5*B-C* and *SI Appendix*, Fig. S5*A-E*). No DDR signaling proteins were recruited to the *LacO* by the negative control, mCherry-LacRnls. As a positive control, we added 4-OHT and Shield-1 to the medium for 6 h, and readily detected ATM, phospho(p)-ATM (Ser1981), γ-H2AX, RAD50, NBS1, and MRE11. No pATM or γ-H2AX signals were detected at the array upon recruitment of mCherry-LacRnls-IE1, consistent with a constitutive interaction between IE1 and NBS1 that is independent of DSB signaling. Intriguingly, while an mCherry-LacRnls-NBS1 fusion protein is sufficient to induce ATM recruitment and activation, as well as subsequent H2AX phosphorylation at the *LacO* array in NIH-3T3 cells (57), NBS1 recruitment by mCherry-LacRnls-IE1 did not trigger ATM activation (Fig. 5B and *SI Appendix*, Fig. S5*B*). To further validate that IE1 inhibits the ability of NBS1 to activate ATM at the *LacO*, we transiently transfected a mCherry-LacRnls-NBS1 fusion protein and an untagged or FLAG-tagged IE1 vector in U2OS 2-6-5 cells. As expected, a full length (FL) NBS1 construct (aa 1–754) specifically promoted ATM and H2AX phosphorylation at the *LacO* in approximately 75% and 50% of cells, respectively (Fig. 5*D-F* and *SI Appendix*, Fig. S5*F-H*). This function depends on its ability to bind ATM, as an NBS1 ATM binding deficient construct (ΔA) (aa 1–733) produced similar γ-H2AX levels as the negative control (Fig. 5*D* and *SI Appendix*, Fig. S5*F*). Interestingly, NBS1-dependent accumulation of γ-H2AX and pATM was strongly inhibited in cells expressing untagged or FLAG-tagged IE1 (Fig. 5*D-F* and *SI Appendix*, Fig. S5*F-H*). These findings suggest that the interaction between IE1 and NBS1 at the array directly prevents ATM activation and subsequent H2AX phosphorylation. In support of this model, we found that IE1 accumulates at the *LacO* array upon DSB induction in an NBS1-dependent manner (Fig. 5*G-H*), an observation that can only be made in this system, as no marker of DSB signaling can be used to detect IE1 accumulation at endogenous DSBs. In these conditions, IE1 colocalizes with 60% of mCherry-LacR-FOKI foci and this amount is reduced to 30% in cells treated with a siRNA against NBS1.

**Fig. 5.**
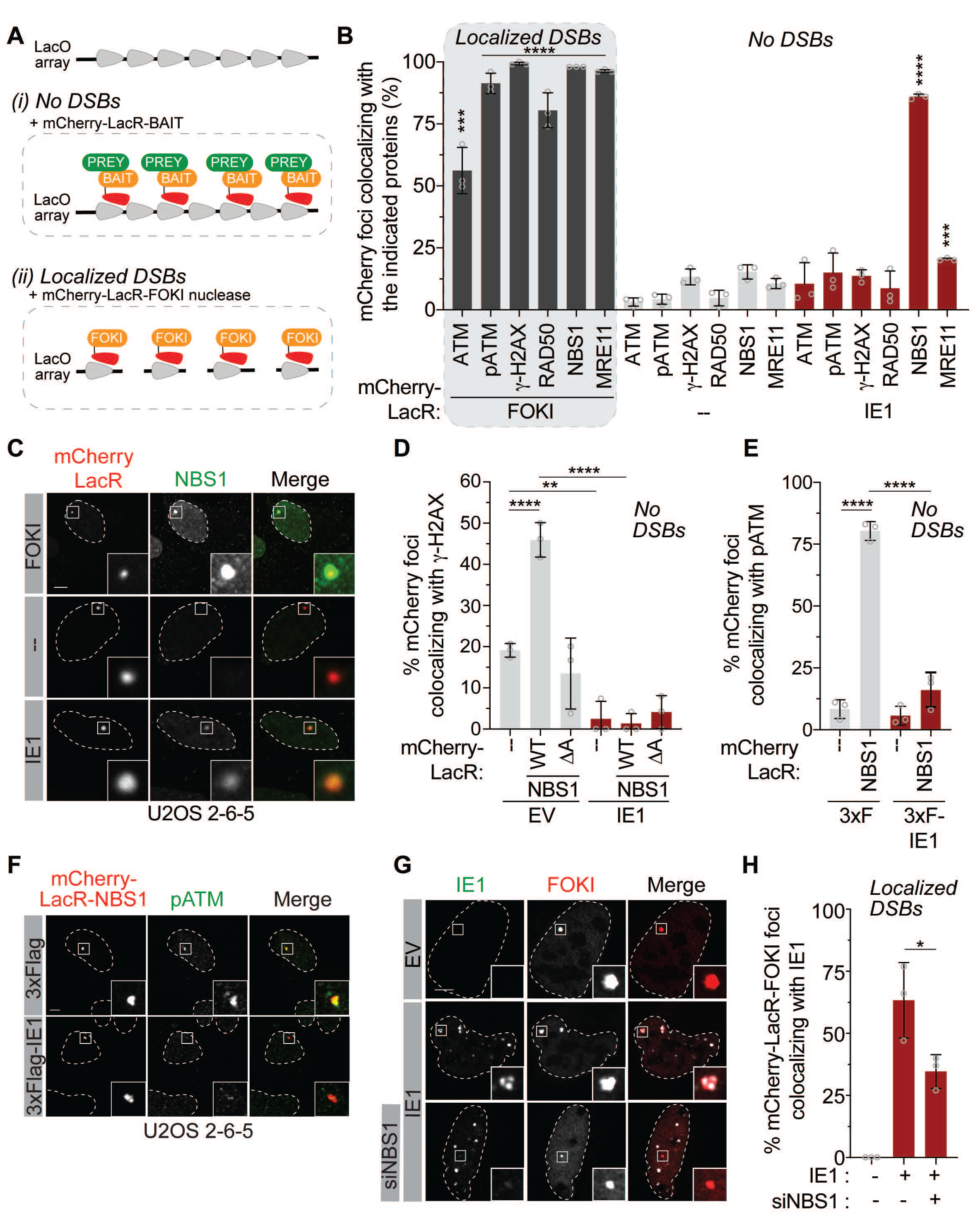
HHV-6B IE1 interacts with NBS1, inhibits ATM activation, and is recruited to DSBs *(A)* The integrated *LacO* array-based assay used to study protein colocalization at a specific locus without DSBs (*i*) or DNA repair protein recruitment at localized DSBs (*ii*). *LacO* array repeats, mCherry-LacRnls-fusion proteins, and preys are shown in grey, red/orange, and green, respectively. *(B)* Quantification of the indicated DSB-signaling proteins at localized DSBs induced by ER-mCherry-LacRnls-FOKI-DD (FOKI) or at either a mCherry-LacRnls (--, negative control) or mCherry-LacRnls HHV-6B IE1 protein foci in U2OS 2-6-5 cells. Transfected cells were treated with 4-OHT and Shield-1 for 6 h then immunostained for ATM, pATM (S1981), γ-H2AX, RAD50, NBS1, and MRE11 (Representative images, *SI Appendix*, Fig. S5*A-E)*. For each condition, statistical significance was analyzed against the control protein (mCherry-LacRnls). Data represent the mean ± SD (*n* = 3). *(C)* Representative images of NBS1 recruitment at DSBs (top panel) and its colocalization with IE1 in the absence of DSBs (bottom panel). *(D-E)* U2OS 2-6-5 cells were treated as described in (B), immunostained for γ-H2AX (D) or pATM (E) (*SI Appendix*, Fig. S5*F-G*) and quantified as indicated. In both experiments, cells were transfected with vectors expressing either untagged or 3×FLAG-IE1. *(F)* Representative images of pATM inhibition at the mCherry-LacRnls-NBS1 locus. Cells were treated as described in (B). Representative images of the negative controls are presented in *SI Appendix*, Fig. S5*G-H. (G)* Representative images of NBS1-dependent IE1 recruitment to DSBs. Cells were treated with a siCTRL or siNBS1 prior to their transfection with an untagged IE1 or an empty vector. ER-mCherry-LacR-FOKI-DD was induced as described in (B). Cells were processed for IE1 immunofluorescence. *(H)* Quantification of the mCherry-LacR FOKI foci colocalizing with IE1. Data represent the mean ± SD (*n* = 3). \*\**p* < 0.05, \*\**p* < 0.01, *** *p*<0.001, \*\*\*\**p* < 0.0001. Scale bars = 5 μm.

### Two distinct domains of IE1 interact with NBS1 and prevent ATM activation

The functional domains of IE1 are not well characterized aside from its STAT2 binding domain (aa 270– 540; Fig. 6*A*) (36). Guided by its secondary structure, we designed a series of IE1 fragments that we fused to mCherry-LacRnls to assess their ability to recruit endogenous NBS1 to the *LacO* array (*SI Appendix*, Fig. S6*A*). The fusion encoding aa 1–540 was the smallest fragment capable of recruiting NBS1 at the *LacO* array as efficiently as FL IE1 (∼81% of mCherry-LacRnls-IE1 1–540 colocalized with NBS1; Fig. 6*B-C* and *SI Appendix*, Fig. S6*B*). All attempts to generate smaller fragments of this N-terminal domain of IE1 resulted in unstable proteins in our hands. Interestingly, the 1–540 fragment was also the smallest to efficiently accumulate in PML bodies (*SI Appendix*, Fig. S6*C-D*) suggesting that both functions are related. Consistently, we observed a reduced accumulation of IE1 at PML bodies in cells treated with siNBS1 (*SI Appendix*, Fig. S4*E-F*).

**Fig. 6.**
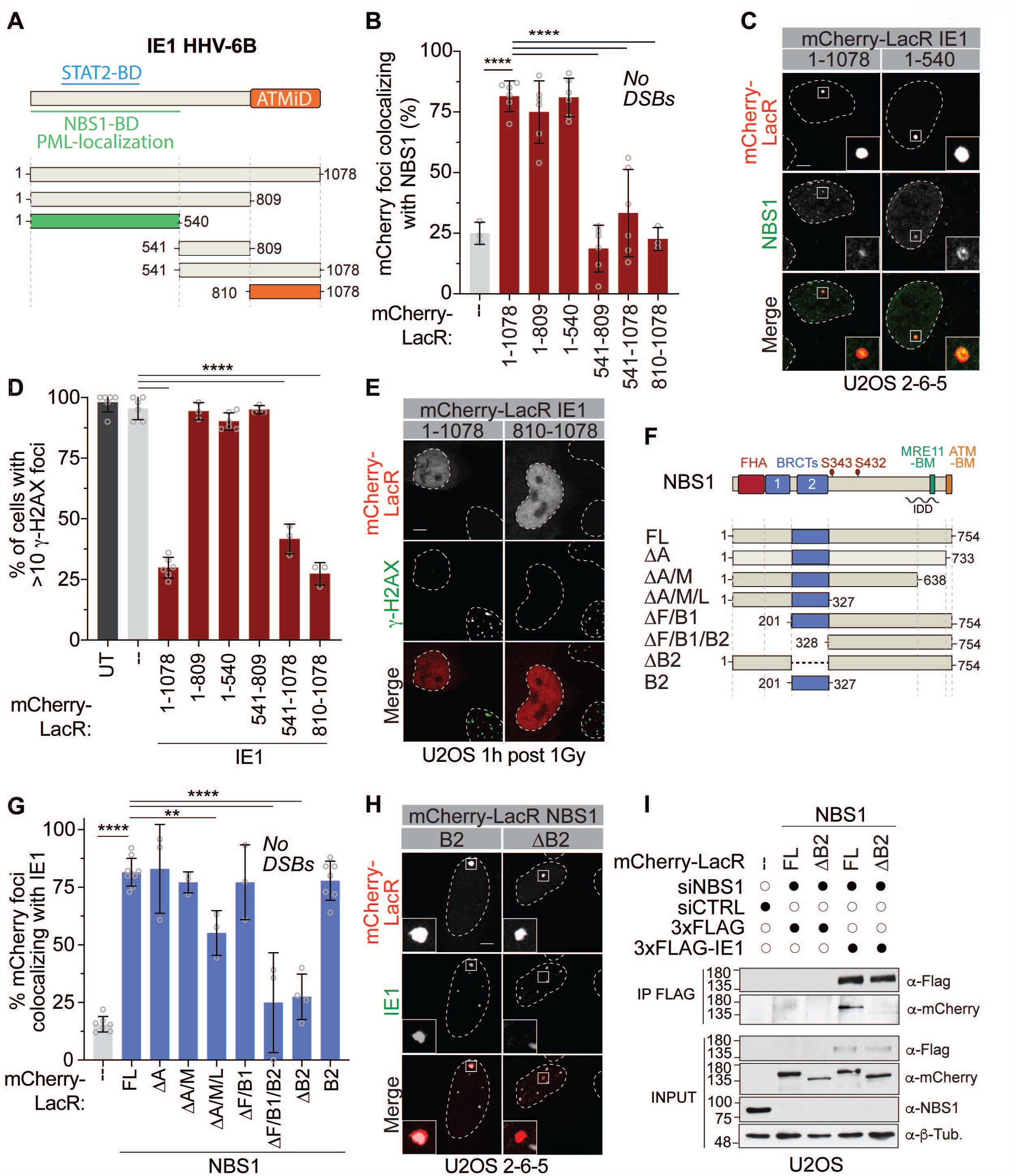
IE1 interacts with NBS1 and inhibits ATM through two distinct domains *(A)* Schematic of HHV-6B IE1 and the protein fragments used in this study. NBS1-BD, NBS1-binding domain; NBS1i, NBS1 inhibitory domain, STAT2-BD: STAT2 binding-domain (aa 270–540). *(B-C)* U2OS 2-6-5 cells transfected with plasmids expressing the indicated mCherry-LacR fusion proteins were immunostained for NBS1 (see also *SI Appendix*, Fig. S6*A-B*). The mCherry-LacR backbone was used as a negative control (--). *(D-E)* Quantification of cells with > 10 γ-H2AX foci. UT, untreated. Transiently transfected cells were irradiated and immunostained for γ-H2AX 1 h later (see also *SI Appendix*, Fig. S6*G*). Untreated cells and the mCherry-LacR backbone were used as negative controls (--). *(F)* Schematic of NBS1 and the protein fragments used in this study. FHA, forkhead-associated domain; BRCT, BRCA1 C-terminal domain; MRE11-BM, MRE11-binding motif; ATM-BM, ATM-binding motif; IDD, intrinsically disordered domain. *(G-H)* U2OS 2-6-5 cells were transfected with the indicated mCherry-LacR plasmids and immunostained for IE1 (*SI Appendix*, Fig. S6*H-M*). *(I)* U2OS cells treated with siCTRL or siNBS1 were transfected with the indicated 3×FLAG and mCherry-LacR constructs. After 24 h, WCEs were prepared and 3×-FLAG-IE1 interactors were immunoprecipitated using anti-Flag (M2) agarose beads and immunoblotted for FLAG, mCherry, and NBS1. β-tubulin (Tub.) was used as a loading control. Data for *(B), (D)*, and *(G)* represent the mean ± SD (*n* = 3). ** *p*<0.01, \*\*\*\**p* < 0.0001. Scale bars = 5 μm.

To determine if the IE1-NBS1 interaction is sufficient to prevent ATM activation, we transiently transfected expression vectors containing the different fragments of IE1 into U2OS cells (without the *LacO* array) and quantified their ability to inhibit H2AX phosphorylation in irradiated cells. Surprisingly, the IE1 N-terminus alone was unable to prevent the accumulation of γ-H2AX foci (Fig. 6*D-E* and *SI Appendix*, Fig. S6*G*). Instead, we found that this function depends on a fragment of 268 amino acids in the IE1 C-terminus. The 810–1078 fragment inhibited H2AX phosphorylation as efficiently as the FL protein (∼75% of cells transfected with mCherry-LacRnls-IE1 810–1078 had <10 γ-H2AX foci 1 h post-irradiation with 1 Gy; Fig. 6*D-E* and *SI Appendix*, Fig. S6*G*). Together, these results show that IE1 interacts with NBS1 and blocks ATM activation using two distinct motifs: an N-terminal NBS1-binding domain (NBS1-BD) and a C-terminal domain that independently inhibits the ability of NBS1 to trigger DSB signaling (Fig. 6*A*), which we have named ATM-inhibitory domain (ATMiD).

NBS1 encodes a 95-kDa protein with multiple domains, which are required for its recruitment to DSBs and its interactions with the ATM and ATR (58). Briefly, NBS1 contains a forkhead-associated (FHA) domain and two breast cancer C-terminal domains (BRCTs) that are required for optimal phospho-dependent NBS1 accumulation at DNA breaks. The C-terminus contains a domain that promotes its interactions with MRE11 (MRE11-binding motif, MBM) and ATM (ATM-binding motif, ATM-BM; Fig. 6*F*). Interestingly, NBS1 also contains an intrinsically disordered domain (IDD) that drives a species-specific interaction with the HSV-1 IE protein ICP0 (59). To determine if this domain also promotes the interaction between NBS1 and IE1, we used the same approach used to map the IE1-NBS1 interaction (Fig. 6*A-C*). Different fragments of NBS1 were fused with the mCherry-LacRnls protein and co-expressed with an untagged version of IE1 (Fig. 6*F-H* and *SI Appendix*, Fig. S6*H-M*). The mCherry-LacRnls-NBS1 construct lacking the BRCT2 domain (ΔB2, Δaa 201–327) was unable to recruit IE1 to the array, while the construct containing only this domain was sufficient for the interaction (Fig. 6*F-H* and *SI Appendix*, Fig. S6*J, M*). Consistently, immunoprecipitation of FLAG-tagged IE1 from U2OS cell lysates revealed an interaction with FL mCherry-LacRnls-NBS1 but not the ΔB2 fusion (Fig. 6*I*). Using the LacR system, we noted that the mCherry-LacRnls-NBS1 fusion lacking the linker region of NBS1 (ΔL, Δ328–638) significantly reduced the interaction between NBS1 and IE1 (*SI Appendix*, Fig. S6*H, J-M*). In contrast with the BRCT2 domain, the linker alone was unable to recruit IE1 to the *LacO* array (*SI Appendix*, Fig. S6*H, J-M*).

Altogether, our results support a model where the N-terminus of IE1 interacts with the BRCT2 domain of NBS1 and the C-terminus of IE1 blocks ATM activation. In the LacR system, IE1 did not interact with the domain of NBS1 that interacts with ATM (ATM-BM, aa 733–754) (Fig. 6*F-H* and *SI Appendix*, Fig. S6*I, L*) or with ATM itself (Fig. 5*B*). The latter observation suggests that IE1 does not interfere with the NBS1-dependent activation of ATM by directly competing for interactions between them or that the interaction is too weak to be detected in our experimental setting.

### HHV-6B integration relies on a pathway that safeguards telomere elongation

Depending on the virus, the MRN complex is either required for viral replication or it inhibits it (7). As HHV-6B IE1 interacts with NBS1 and blocks ATM activation, the complex is likely detrimental for its replication. HHV-6B infection has different outcomes depending on the nature of the infected cells (Fig. 7*A*). In permissive cells (*e*.*g*., MOLT-3), viral protein expression promotes replication (the lytic state). In contrast, in semi-permissive cells, integration of the viral genome into the host’s telomeres is favored, and this process has been proposed to rely on HDR processes in the infected cells (24). To understand the interplay between HHV-6B, DSB signaling, and HDR repair, we investigated the impacts of depleting NBS1 on viral replication and integration. In these experiments, we depleted NBS1 from permissive cells (MOLT-3) and semi-permissive cells (U2OS, HeLa, and GM847) by shRNA (*SI Appendix*, Fig. S6*A-D*). MOLT-3 cells treated with control and NBS1 shRNA were infected with HHV-6B and viral DNA was quantified over time by qualitative polymerase chain reaction (qPCR; Fig. 7*B*). Viral DNA replication was 1.67-fold higher in MOLT-3 cells depleted of NBS1 72 h post-infection (note that this is an underestimate, as CellTiter-Glo® assays revealed that the shRNA against NBS1 was toxic in MOLT-3 cells, Fig. 7*C*). Viral integration was assessed in two types of semi-permissive cells: HeLa cells, which lengthen their telomeres *via* hTERT-dependent mechanisms, and U2OS and GM847 cells which rely on ALT, a telomerase-independent mechanism that uses HDR pathways for telomere elongation (60, 61). All cell lines were infected with HHV-6B at a multiplicity of infection (MOI) of 1 and passaged for 4 weeks prior to DNA extraction and viral genome quantification by droplet digital (dd)PCR (62). Interestingly, levels of viral integration were approximately 6-fold higher in HeLa cells depleted of NBS1 than in control HeLa cells (Table 1). In contrast, the integration frequency was decreased by at least 2-fold in U2OS and GM847 cell lines depleted of NBS1 vs the control lines. This difference resembles the lower integration level measured in U2OS *PML*^−/−^ cells, a condition previously reported to reduce viral integration (Table 1) (42). NBS1 depletion did not further impact viral integration in these conditions. Lastly, the differences in viral integration levels between the semi-permissive cell lines were not artefactually driven by cell death, as shNBS1 slightly decreased the viability of all cell types used in this study (Fig. 7*C* and *SI Appendix*, Fig. S7E). Altogether, these results are consistent with the need for functional NBS1-dependent HDR repair pathways to promote integration in ALT^+^ cells and support a model where viral integration in semi-permissive cells relies on the molecular mechanisms that drive telomere elongation rather than specific DNA repair mechanisms.

**Fig. 7.**
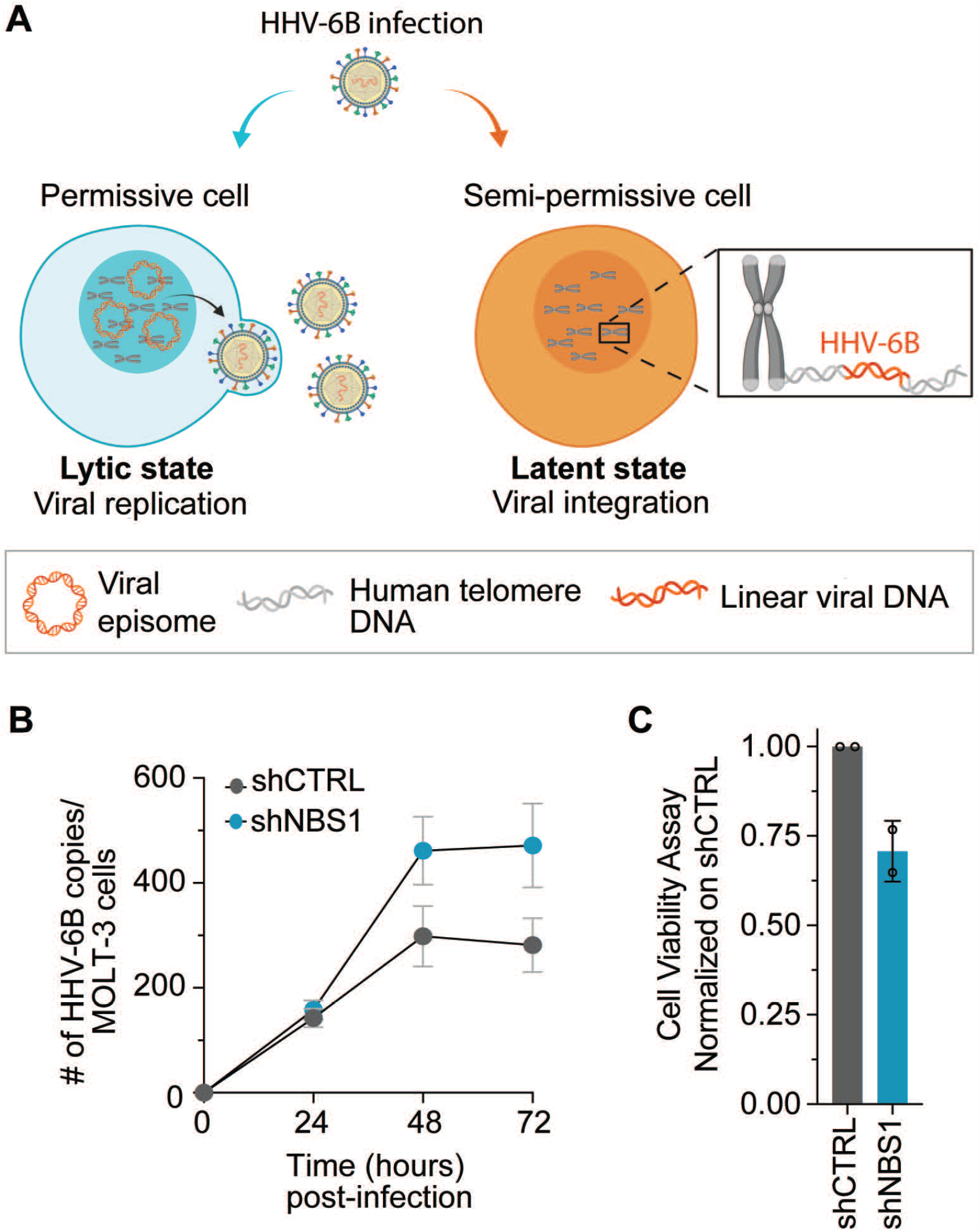
NBS1 depletion impairs viral integration in cells maintaining their telomeres by homology-directed repair HHV-6B infection in permissive and semi-permissive cells. In cells semi-permissive for HHV-6B, replication is inefficient, and the viral genome integrates at the telomeres. *(B)* MOLT-3 cells with and without NBS1 (*SI Appendix*, Fig. S7*A*) were infected with HHV-6B at a MOI of 1 and harvested at the indicated time points. Following cell lysis, DNA was extracted and HHV-6B was quantified by qPCR using primers for HHV-6B *U67-68* and human *RPP30*. Data represent the mean ± SD (*n* = 3). *(C)* MOLT-3 cells were transduced with the indicated shRNA and passaged 5 times prior to CellTiter-Glo® analyses. Cell viability was determined using standard curves for each cell line and normalized to the shCTRL condition for each experiment.

**Table 1.** Importance of NBS1 for HHV-6B chromosomal integration in ALT^+/−^ cells

| Cell line | ALT status | shRNA | % cells with integrated HHV-6B<sup>a</sup> (n)<sup>b</sup> | *P* value<sup>c</sup> |
| --- | --- | --- | --- | --- |
| HeLa | Negative | CTRL | 0.96 (36,320) | <2.2e<sup>-16</sup> |
|  |  | NBS1 | 6.11 (33,280) |  |
| GM847 | Positive | CTRL | 0.65 (21,820) | <2.2e<sup>-16</sup> |
|  |  | NBS1 | 0.01 (18,320) |  |
| U2OS | Positive | CTRL | 1.60 (20,000) | <2.2e<sup>-16</sup> |
|  |  | NBS1 | 0.69 (21,520) |  |
| U2OS *PML*<sup>-/-</sup> | Positive | CTRL | 0.71 (28,220) | ns<sup>d</sup> |
|  |  | NBS1 | 0.78 (30,460) |  |

<sup>a</sup> mean of three independent cultures<sup>b</sup> total number of cells analyzed<sup>c</sup> Pearson's Chi-squared test with Yates' continuity correction<sup>d</sup> ns, not significant

## Discussion

In this study, we sought to understand how HHV-6B manipulates factors that safeguard genomic instability in infected cells, as well as its impacts on two key events of the viral life cycle: genome replication and chromosomal integration. Using a series of microscopy- and cytometry-based approaches to track the source of DNA breaks in infected cells and in cells expressing the HHV-6B immediate-early protein IE1, we found that IE1 promotes the accumulation of DNA DSBs and inhibits their repair. Further structure-function analyses revealed a molecular mechanism by which HHV-6B IE1 localizes to DSBs in an NBS1-dependent manner and prevents HDR-mediated DNA repair by blocking ATM activation and subsequent DDR signaling. We report that IE1 specifically interacts with NBS1 through an N-terminal NBS1-BD and that ATM activation by NBS1 is strongly inhibited by a newly characterized domain of IE1, the C-terminal ATMiD.

ATM activation by the MRN complex requires conformational changes in ATM that expose its substrate-binding site (53). Our findings show that, in contrast with NBS1, IE1 does not interact strongly with ATM in the LacR-based system. Furthermore, the ATMiD domain of IE1 does not interact with NBS1. The exact mechanism by which IE1 inhibits ATM activation thus remains unclear. Based on current literature, IE1 could be interfering with ATM activation by preventing the interaction between the FxF/Y motif of NBS1 and ATM or by directly blocking the substrate-binding site of ATM (53, 63). Alternatively, IE1 may block H2AX phosphorylation through steric hindrance (*e*.*g*., though chromatin binding). Viral proteins such as Kaposi’s sarcoma-associated herpesvirus LANA and adenovirus protein VII interfere with the activation of chromatin-dependent mechanisms by directly interacting with the nucleosomes (64, 65). Further investigation will be required to elucidate how IE1 prevents ATM activation by the MRN complex.

The mechanism by which IE1 interacts with NBS1 and inhibits ATM signaling in cells differs from the mechanisms by which other viruses manipulate this pathway (2, 59). The BRCT2 domain of NBS1 contributes to its retention on DSBs (66), which may be reduced when IE1 binds this domain. However, the fact that the IE1 N-terminus is insufficient to inhibit ATM signaling suggests otherwise. In this study, we show that IE1 is recruited to DSBs in an NBS1-dependent manner and that ectopic expression of the IE1 ATMiD is sufficient to inhibit DSB signaling. Thus, it is still unclear whether IE1 needs to accumulate at DSBs in an NBS1-dependent manner to inhibit ATM when expressed in infected cells at lower levels, or if NBS1 interaction and ATM inhibition are independent functions of IE1. A model in which IE1 inhibits ATM activation through a bi-partite mechanism is appealing, as it would provide a way for HHV-6B to inhibit ATM signaling at specific loci. This would support a recently proposed concept in which viruses prevent local ATM signaling on the viral genome and restrict viral replication, while avoiding a global inhibition of the DSB signaling cascade in infected cells (67). During the lytic cycle, the accumulation of genomic instability in the host cell genome is not a problem as these cells will die upon the lysis provoked by the virus to release new virus particles. However, more selective inhibition of ATM by IE1 during the latent cycle of HHV-6B or from iciHHV-6B would avoid a detrimental accumulation of genomic alterations in host cells. This model would be consistent with the fact that HHV-6B is not associated with a higher frequency of cancer development, as would be expected if global DSB signaling was inhibited in these cells. Alternatively, expression of IE1 upon the exit of latency may inhibit global DSB signaling, but this phenomenon is restricted to the early stages of the process, thereby minimizing the impact on the host cell’s genomic stability.

In addition to its role during viral infection, Peddu et al. used RNA-seq approach to show that *IE1 (U90)* is among the most abundantly expressed genes in a variety of tissues from iciHHV-6A/B+ individuals (68). Spontaneous and inducible IE1 protein expression from integrated HHV-6A/B genomes has also been documented (62), raising the possibility that IE1 expression from integrated genomes might contribute to the development of iciHHV-6A/B associated diseases by inducing genomic instability in these cells. At present, conditions associated with iciHHV-6A/B status include increased spontaneous abortion rates (27, 28), pre-eclampsia (29) and angina pectoris (30). Further characterization of the proteins expressed from integrated genomes as well as the diseases associated with these conditions will be required to strengthen our understanding of the consequences associated with viral latency in iciHHV-6A/B subjects. Importantly, the intricate interplay between IE1, the MRN complex, and ATM pathway activation will need to be studied in a spatiotemporal manner to elucidate when and how IE1 manipulates this important pathway during viral infection and integration. Further efforts will also be required to determine if ATM inhibition by IE1 contributes to its ability to block type I interferon signaling in infected cells (36). From a mechanistic point of view, it will be interesting to investigate if the interaction between IE1 and NBS1’s BRCT2 domain—a phospho-recognition domain—is regulated by phosphorylation (66, 69–72). Finally, the model presented here assumes that NBS1 and ATM activity must be inhibited to prevent their detrimental effect on viral replication. However, it is impossible to rule out that enhanced viral replication and integration result from the increased level of genomic instability induced in host cells upon viral infection. Further studies will be required to address this question.

In germline, hematopoietic, stem, and rapidly renewing cells, telomere elongation relies on hTERT, a polymerase that catalyzes the extension of telomeric DNA repeats using RNA as a template (73). While hTERT is negatively regulated in healthy somatic cells, cancer cells can overcome senescence either through its re-activation or by an alternative homology-directed mechanism called ALT(60). The HHV-6B genome contains conserved telomeric sequences that are required for viral integration (23). In this study, we show that HHV-6B integration is independent of NBS1 in ALT^−^ cells but dependent on NBS1 in ALT^+^ cells. These findings are consistent with previous reports showing that the telomerase complex is required for optimal HHV-6B integration (74) and with the reported role of NBS1 in ALT (75, 76). While PML is not required for the IE1-NBS1 interaction or the ability of IE1 to inhibit H2AX phosphorylation (this study), NBS1 is required for the assembly of functional ALT-associated PML bodies (77). These concomitant roles are in line with the absence of phenotypes associated with NBS1 depletion in integration assays performed on *PML*^−/−^ ALT^+^ U2OS cells. Intriguingly, we previously reported that *PML* knockout also reduces integration in ALT^−^ HeLa cells, reinforcing the hypothesis that PML plays an ALT-independent role in this process (42). Further studies will be required to elucidate this function.

Consistent with previous findings showing that HHV-6B integration is not altered upon inhibition of RAD51 (78, 79), we found that IE1 inhibits HDR processes, and that integration is independent of NBS1 in ALT-cell lines. Together, these observations argue against models where integration mechanisms rely on RAD51-dependent BIR or SSA (24). However, it is important to note that all homology-directed reporter assays used in this study rely on extensive DNA end resection following nuclease-induced breakage, a process that is dependent on NBS1 (80). Thereby, integration models where SSA or RAD51-independent BIR trigger integration following extensive accumulation of ssDNA generated at stalled replication forks remain plausible. One attractive model is that HHV-6B integration occurs during mitotic DNA synthesis (MiDAS), a RAD52-dependent BIR mechanism that is initiated by replication fork stalls that remain unresolved at the start of mitosis—a problem often observed at hard-to-replicate loci, including the telomeres (13, 27, 81). Alternatively, upon cell entry but before viral genome circularization (and before IE1 is expressed), the viral genome may be perceived as broken DNA. Under such circumstances, the MRN complex would be recruited to the ends of the viral genome and initiate 3′→ 5′ resections. The ssDNA ends of eroded telomeres (no longer efficiently protected by the shelterin complex) could anneal to the near-terminal telomeric sequence at the right end of the genome in a process analogous to an ALT mechanism described in yeast (reviewed in (13)). Once the entire viral genome is copied, the telomeric repeats at the left end of the genome would serve as a template for the hTERT and ALT mechanisms to regenerate a telomere of appropriate length (82).

In conclusion, we provide a detailed characterization of the HHV-6B IE1 protein as an efficient inhibitor of DSB signaling and DDR that contributes to the favorable establishment of a productive infection. Despite being a relatively abundant protein expressed very early upon entry, the functions of IE1 remain poorly defined. IE1 shares very little sequence homology with proteins from other herpesviruses (except HHV-6A and HHV-7) meaning that deductions based on primary sequence analysis are very limited. Our work adds to the growing knowledge surrounding HHV-6B integration processes and the potential importance of IE1 during infection.

## Materials and Methods

### RNA interference

A SMARTPool siRNA targeting RAD51, single siRNA duplexes targeting NBS1, and a non-targeting single siRNA duplex were purchased from Dharmacon (Horizon). Single siRNA duplexes targeting RAD52 were a kind gift from Jean-Yves Masson (Université Laval, Québec, Canada). siRNAs were forward-transfected 24 h prior to cell processing using RNAimax (Invitrogen) according to the manufacturer’s protocol. Plasmids carrying an NBS1 shRNA (Open Biosystems) or a control shRNA (Sigma) in the pLKO background backbone were a kind gift from Cary A. Moody (6). Lentiviruses were produced as previously described (6). Briefly, plasmids expressing shRNAs with vesicular stomatitis virus G (pMD2.g) and lentiviral packaging (pPAX) plasmids were co-transfected into HEK293T cells using polyethyleneimine. Lentivirus-containing supernatants were harvested 48–72 h post-transfection, and U2OS, MOLT-3, HeLa, and GM847 cells were transduced in the presence of 8 μg/mL Polybrene (Sigma). For all relevant experiments, RAD51, RAD52, and NBS1 depletion was confirmed by immunoblotting or qPCR analyses.

### Cell cultures and transfections

Cell lines were maintained at 37°C and 5% CO_2_. All culture media were supplemented with 10% fetal bovine serum. MOLT-3 cells (American Type Culture Collection (ATCC)) were cultured in Roswell Park Memorial Institute-1640 (RPMI) medium (Corning Cellgro), 8.85 mM HEPES, and 5 μg/mL plasmocin (Invivogen). GM847 and HeLa cell lines were obtained from ATCC and cultured in Dulbecco’s modified Eagle’s medium (DMEM; Corning Cellgro), NEM (Corning Cellgro), 8.85 mM HEPES, and 5 μg/mL plasmocin. U2OS (ATCC), U2OS *PML*^−/−^ (42), U2OS 2-6-5 (a kind gift from Roger Greenberg, University of Pennsylvania, Philadelphia) (55), U2OS DR-GFP, SA-GFP (a kind gift from Jeremy Stark, City of Hope National Medical Center, California) (48, 52), and cell lines were cultured in McCoy’s medium (Life Technologies).

### Viral infection and integration assays

Viral infection was done as previously described (42) using HHV-6B (strain Z29) at an MOI of 1 (or not (Mock samples)). At the indicated time points, cells were harvested and processed for DNA extraction using a QIAamp DNA Blood Mini Kit (Qiagen) and analyzed by qPCR. Integration assays were performed as described previously (62). Briefly, cells were infected with HHV-6B (MOI of 1) for 24 h and passaged for 4 weeks prior to DNA extraction with the QIAamp DNA Blood Mini Kit for ddPCR.

### PCR analyses

qPCR was performed as previously described (83). DNA was quantified using primers and probes against *U67-68* (HHV-6B) and ribonuclease P/MRP subunit p30 (*RPP30*; as a host reference gene). Data were normalized against the corresponding genome copies of *RPP30*. ddPCR was used to quantify integration frequency as previously described (62). Briefly, HHV-6B chromosomal integration frequencies were estimated assuming a single integrated HHV-6/cell and calculated with the following formula: (number of HHV-6 copies)/(number of *RPP30* copies/2 copies per cell) × 100, as previously described (62). This assay has been extensively validated and provides comparable data to single cell cloning and quantification.

### Immunofluorescence microscopy

Immunofluorescence were done essentially as previously described for MOLT-3 (42) and U2OS (56) cells. Briefly, cells were either fixed with 2% paraformaldehyde (PFA) or 100% MeOH prior to permeabilization and incubation with primary antibody diluted in blocking buffer. DNA was counterstained with DAPI and the coverslips were mounted onto glass slides with Prolong Diamond Mounting Agent (Invitrogen). Further experimental details are provided as the Supplementary information.

### FISH

Fixed cells were processed as described for immunofluorescence staining and then fixed for 2 min at room temperature with 1% PFA/PBS. Coverslips were washed twice with PBS for 5 min and dehydrated for 5 min in successive ethanol baths (70%, 95%, 100%). Once dried, coverslips were placed upside down on a drop of hybridizing solution (70% formamide, 0.5% blocking reagent (Sigma, Cat:11096176001), 10 mM Tris-HCl pH 7.2, 1/1000 Cy5-TelC PNA probe (F1003, PNABio)). Samples were denatured for 10 min at 80°C on a heated block, then incubated overnight at 4°C in the dark. After hybridization, coverslips were washed twice for 15 min in washing solution (70% formamide; 10 mM Tris-HCl pH 7.2) and then washed three times for 5 min with PBS. Coverslips were air-dried, counterstained with DAPI, washed with PBS, and mounted onto glass slides with Prolong Gold Mounting Agent.

### Metaphase spread analysis

U2OS SA-GFP HHV-6B IE1 cells were arrested in mitosis using 1 μM nocodazole for 3 h at 37°C and 5% CO_2_. Cells were then resuspended and incubated in pre-warmed hypotonic solution (KCl 75 mM, 15% fetal bovine serum) at 37°C for 15 min to induce swelling and fixed in a 75% ethanol 25% acetic acid solution overnight at 4°C. Droplets of cells were spread onto glass slides pre-cooled to -20°C and dried overnight in the dark at room temperature. Slides were then mounted with Vectashield Antifade Mounting Medium containing DAPI (VECTH20002, MJS BioLynx Inc.). Images were acquired using a Zeiss LSM700 laser-scanning microscope equipped with a 40× water lens. Quantification was performed on three biological replicates and 10 spreads were quantified per experiment.

### Immunoprecipitation

U2OS cells (1×10^7^) were transfected with NBS1- or non-targeting single siRNA duplexes for 24 h, then co-transfected with the indicated mCherry-LacR and 3×FLAG expression vectors. After 24 h, cells were lysed in NETN lysis buffer (50 mM Tris pH 8.0, 150 mM NaCl, 1mM EDTA, 0.5% NP-40) complemented with 1× complete, EDTA-free Protease Inhibitor Cocktail (Roche), 20 mM N-ethylmethylamine, 1 mM NaF, and 0.2 mM Na_3_VO_4_. Cleared cell lysates were immunoprecipitated using 1 μg FLAG-M2 antibody coupled to 40 μL of packed protein G Sepharose beads (Cat GE17-0618-01, Sigma) for 3 h at 4°C. Beads were washed four times with NETN buffer and eluted in 2× Laemmli buffer for immunoblotting.

### Reporter-based DNA repair assays

DR-GFP, NHEJ-GFP, SA-GFP, and BIR-GFP cell lines were plated at 125,000 cells/well in 6-well plates. After 24 h, cells were co-transfected with 900 ng of the I-SceI plasmid (pCBASceI, Addgene #26477) and 900 ng of pcDNA4/TO-HHV-6B IE1 (+I-SceI, +IE1) or 900 ng of the pcDNA4/TO/Myc-His vector as a negative control (+I-SceI,-IE1). The pcDNA4/TO/myc-His vector alone was transfected for conditions without IE1 and I-SceI (-I-SceI/-IE1). A plasmid expressing iRFP (200 ng) was also transfected into all conditions to control for transfection efficiency. After 48 h, cells were harvested and washed with PBS, and an Accuri C6 flow cytometer (BD Biosciences) was used to quantify the GFP^+^ cells in the iRFP^+^ population. Data were analyzed using FlowJo. The NHEJ-GFP (EJ7) assay was performed essentially as described above, but cells were co-transfected with 600 ng of each Cas9/sgRNA-expressing vector p330X-sgRNA7a, and p330X-sgRNA7b along with 600 ng of pcDNA4/TO-HHV-6B IE1 or pcDNA4/TO/myc-His (52) and processed for flow cytometry analysis 72 h post-transfection.

### Statistical analysis

Quantifications were performed on three biological replicates. Unless otherwise stated, one-way analysis of variance and Dunnett’s multiple comparisons test were used to assess statistical significance.

## Acknowledgements

We are grateful to Matthew D. Weitzman, Alexandre Orthwein, Cary A. Moody, and members of the Fradet-Turcotte and Flamand laboratories for critical reading of the manuscript; High-Fidelity Science Communications for manuscript editing, and Daniel Durocher, Jean-Yves Masson, Graham Dellaire, Roger Greenberg, and Jeremy Stark for essential reagents. E.B, and V.C. received postdoctoral and doctoral fellowships, respectively, from the Fonds de Recherche du Québec - Santé. V.T. received a master’s fellowship from the Fonds de recherche Nature et technologies. This work was supported by three Canadian Institutes of Health Research Grants (PJT_152948 to A.F.-T.; MOP_123214 and PJT_156118 to L.F.). A.F.-T. is a Tier 2 Canada Research Chair in Molecular Virology and Genomic Instability and is supported by the Foundation J.-Louis Lévesque. We thank the Bioimaging platform of the Infectious Disease Research Centre, which is funded by an equipment and infrastructure grant from the Canadian Foundation for Innovation.

