## Supplemental material for "The immediate early protein 1 of human herpesvirus 6B counteracts ATM activation in an NBS1-dependent manner"

­

Amélie Fradet-Turcotte

Louis Flamand

**This PDF file includes:**

Supplementary Information materials and methods

Figures S1 to S7

Tables S1 to S3

SI References

**Supplementary Information materials and methods**

**Plasmids and virus**

pcDNA4/TO/myc-His-HHV-6B IE1 was previously described (34). The PiggyBac transposon-based (PB)-TetO and PB-CA-rtTA-IRES-NEO plasmids were generated as previously described (82). PB-TetO-HHV-6B IE1, mCherry-LacR, and GFP expression vectors were generated using Gateway^T^ recombination cloning (Invitrogen) and the following destination vectors: pDEST-PB-TetO (82), pDEST-mCherry-LacRnls (83), pDEST-FRT-TO-3×FLAG (ref), and pDEST-FRT-TO-eGFP-NLS (84). HHV-6B IE1 was PCR-amplified from pcDNA4/TO-HHV-6B IE1. HHV-6B IE1 fragments (aa 1–1078, 1–809, 1–540, 541–809, 541–1078, 810–1078) and NBS1 fragments (1–754, 1–733, 1–638, 1–327, 109–754, 201–754, 328–754, ∆201–638, ∆201–327, ∆328–638, 201–327, 328–638, 201–638) were PCR-amplified from pcDNA4/TO-HHV-6B IE1 and pLXIN2-NBS1 (a kind gift from Cary A. Moody, University of North Carolina at Chapel Hill, North-Carolina) (6). HHV-6B strain Z29 (85) was produced in our laboratory as previously described (86). A list of plasmids used in this study is provided in Table S1.

**Cell cultures and transfections**

Dox-inducible U2OS HHV-6B IE1 clones C10 and C102 were established by co-transfecting PB-TetO-HHV-6B IE1, pCMV-hypBAse, and PB-CA-rtTA-IRES-NEO plasmids at a DNA ratio of 1:1:1 into the U2OS SA-GFP cell line using Lipofectamine 2000 (Invitrogen) according to the manufacturer’s protocol. Clones were selected using 40 mg/mL of G418 and isolated by limited dilution. U2OS BIR cells were established by transfecting a GFP-based reporter plasmid (pBIR-GFP) containing an already characterized I-SceI reporter cassette to monitor BIR (49) using Lipofectamine 2000 according to the manufacturer’s protocol. Clones were selected using 2 µg/mL puromycin and isolated by limited dilution. Experiments were performed with a stable reporter clone that produced 1.5–3% GFP^+^ cells after DSB induction. Unless otherwise indicated, IE1 expression was induced for 48 h with 1 µg/mL Dox. HeLa DR-GFP (a kind gift from Roger Greenberg), NHEJ-GFP (EJ7; a kind gift from Jeremy Stark, City of Hope National Medical Center, California), and HEK293T cell lines were cultured in DMEM. All cell lines were validated using short tandem repeat markers and were negative for mycoplasma contamination.

**Chemicals and sources of DNA damage**

Dox (1 µg/mL, D3447, Sigma) was used to induce HHV-6B IE1 expression in stable U2OS cell lines for 48 h. In the FOKI system, DSBs were created at the *LacO* array by promoting the nuclear localization (4-OHT, 100 nM, #3412, Tocris) and stabilization (Shield-1 ligand, 0.5 µM, CIP-S1-0001, CheminPharma) of mCherry-LacR-FOKI nuclease fused to a destabilization domain (DD) and to a modified estradiol receptor (ER) (ER-mCherry-LacR-FOKI-DD) for 6 h prior to fixation for immunofluorescence. DNA damage was also induced by exposing U2OS cells to 1 Gy IR with a CellRad (Precision X-Ray Inc.) and MOLT-3 cells to 4 Gy with a Gammacell^®^ 40 Exactor (Best Theratronics Ltd.).

**RNA extraction and RT-qPCR**

Total RNA was extracted with the RNeasy Mini Kit according to the manufacturer’s instructions (Qiagen) and quantified on a NanoDrop spectrophotometer. Total RNA (250–500 ng) was reverse transcribed with the High-Capacity cDNA Reverse Transcription Kit (Invitrogen) according to the manufacturer’s instructions. Contaminant genomic DNA was removed with a DNaseI (1 unit; ThermoFisher) incubation at 37°C for 2 h prior to the RT reaction and confirmed by performing glyceraldehyde 3-phosphate dehydrogenase (GAPDH) RT-PCR on DNaseI-treated reactions. qPCR was performed for 40 cycles of 94°C for 15 s, 56°C for 5 s, and 72°C for 15 s using a LightCycler 480 apparatus (Roche), LightCycler 480 SYBR Green 1 qPCR Master Mix (Roche), and 5% of the RT reaction as the template. A standard curve was generated by serial dilution, using the U2OS cDNA as the template. The relative expression of each gene was determined using the standard curve and normalized to GADPH. All primer sequences are listed in Table S2.

**Immunofluorescence microscopy**

Irradiated MOLT-3 cells were incubated for 1 h, then pelleted at 900 x g for 5 min and washed three times in PBS. Then, 1×10^4^ cells were added to each well of a microscope slide with 10 reaction wells (MP Biomedicals™ Multitest Slides, Fisher Scientific #ICN6041805). Once dried, cells were fixed for 20 min at room temperature with 2% paraformaldehyde (PFA), hydrated for 5 min with PBS, and processed for immunofluorescence. U2OS and U2OS 2-6-5 cells were grown in 24-well plates on glass coverslips and fixed 24 h later with either 2% (wt/vol) PFA in PBS for 20 min at room temperature or 100% MeOH for 20 min at -20°C. When indicated, cells were irradiated for 15 min or 1 h prior to fixation. For immunostaining with NBS1 and MRE11 antibodies, nuclear extraction with ice-cold NuEx buffer (20 mM HEPES pH 7.4, 20 mM NaCl, 5 mM MgCl_2_, 0.5% NP-40, Complete EDTA-free Protease Inhibitor (Sigma), and 1 mM dithiothreitol) was performed for 20 min on ice prior to fixation. PFA-fixed U2OS cells were further permeabilized with 0.3% (vol/vol) Triton X-100 for 20 min at room temperature. Fixed MOLT-3 and U2OS cells were incubated with blocking buffer (2% bovine serum albumin (BSA) in PBS or 0.1% BSA, 3% goat serum, 0.1% Triton X-100, and 1 mM EDTA pH 8.0 in PBS) for 30 min at room temperature, then with primary antibodies (Table S3) diluted in blocking buffer for 2 h at room temperature, then washed with PBS. Next, cells were incubated with secondary antibodies diluted in blocking buffer for 1 h at room temperature and counterstained with 4',6-diamidino-2-phenylindole (DAPI, 0.4 μg/mL) in PBS. Cells were washed with 1× PBS and the coverslips were mounted onto glass slides with Prolong Diamond Mounting Agent (Invitrogen). To visualize micronuclei in infected and mock MOLT-3 cells, cells were centrifuged at 900 x g for 5 min, resuspended with 10 mL of hypotonic solution (75 mM KCl) for 20 min at 37°C and centrifuged at 900 x g for 5 min. Cell pellets were then resuspended and fixed with fresh 3:1 methanol:acetic acid for 5 min. Cell pellets were washed three times with 3:1 methanol:acetic acid and centrifuged at 900 x g for 5 min. Washed pellets were resuspended in 500 µL of 3:1 methanol:acetic acid, dropped on microscope slides, and air dried prior to DNA counterstaining with DAPI. Images were either acquired using a Zeiss LSM700 (or a recently acquired LSM900) laser-scanning microscope equipped with a 63× oil lens or a Wave FX-Borealis-Leica DMI 6000B microscope with Image EM (Hamamatsu, 512×512 pixels) and Orca-R2 (Hamamatsu, 1344×1024 pixels) cameras and a 40× lens (Quorum Technologies). Images were analyzed and quantified using ImageJ (National Institutes of Health). In micrographs, dashed lines indicated nucleus outlines when DAPI is not shown. Unless otherwise stated, insets represent 10× magnifications of the indicated fields.

**Cell cycle analyses**

Cell cycle analyses were performed as previously described (54, 55). Briefly, cells were fixed with 70% ethanol overnight at 4°C, washed with PBS, and stained with 0.02 mg/mL propidium iodide (PI) in PBS containing 0.1% Triton-X-100 and 0.2 mg/mL RNAse A for 30 min at room temperature. PI staining was detected by flow cytometry (10,000 cells/sample) and cell cycle analyses were performed using FlowJo software (Flow Jo LLC).

**Cell viability assays**

About 15000 cells subjected to NBS1 or control RNAi were added to a 96-well plate and cell viability was evaluated by CellTiter-Glo® Cell Viability Assay (Promega) according to the manufacturer’s instructions. The results were quantified using an Infinite M1000 PRO (Tecan) reader. Standard curves were generated for each cell line to evaluate their viability.

**Reporter-based DNA repair assays**

Validation of BIR-GFP assay. Cells were transfected with RAD51, RAD52, or control siRNA 24 h prior to transfection with the I-SceI expression plasmid. For DNA repair assays in U2OS stable cell lines containing a Dox-inducible expression cassette for HHV-6B IE1 (the SA-GFP and CRISPR-LMNA HDR assays), cells were seeded at 10,000 cells/well in 24-well plates (CRISPR-LMNA-HDR) or at 125,000 cells/wells in 6 well plates (SA-GFP) and induced with 1 µg/mL Dox 24 h prior transfection. The CRISPR-LMNA HDR assay was performed by transfecting 300 ng of the Cas9/*LMNA* sgRNA expression plasmid (pX330-LMNAgRNA1), and 300 ng of CR2.1-mRuby-2-LMNA-Donor in U2OS cells (48). mRuby^+^ cells were analyzed by microscopy using a Ti-LAPP total internal reflection fluorescence microscope (Nikon) 48h post-transfection.

**Figures S1 to S7**

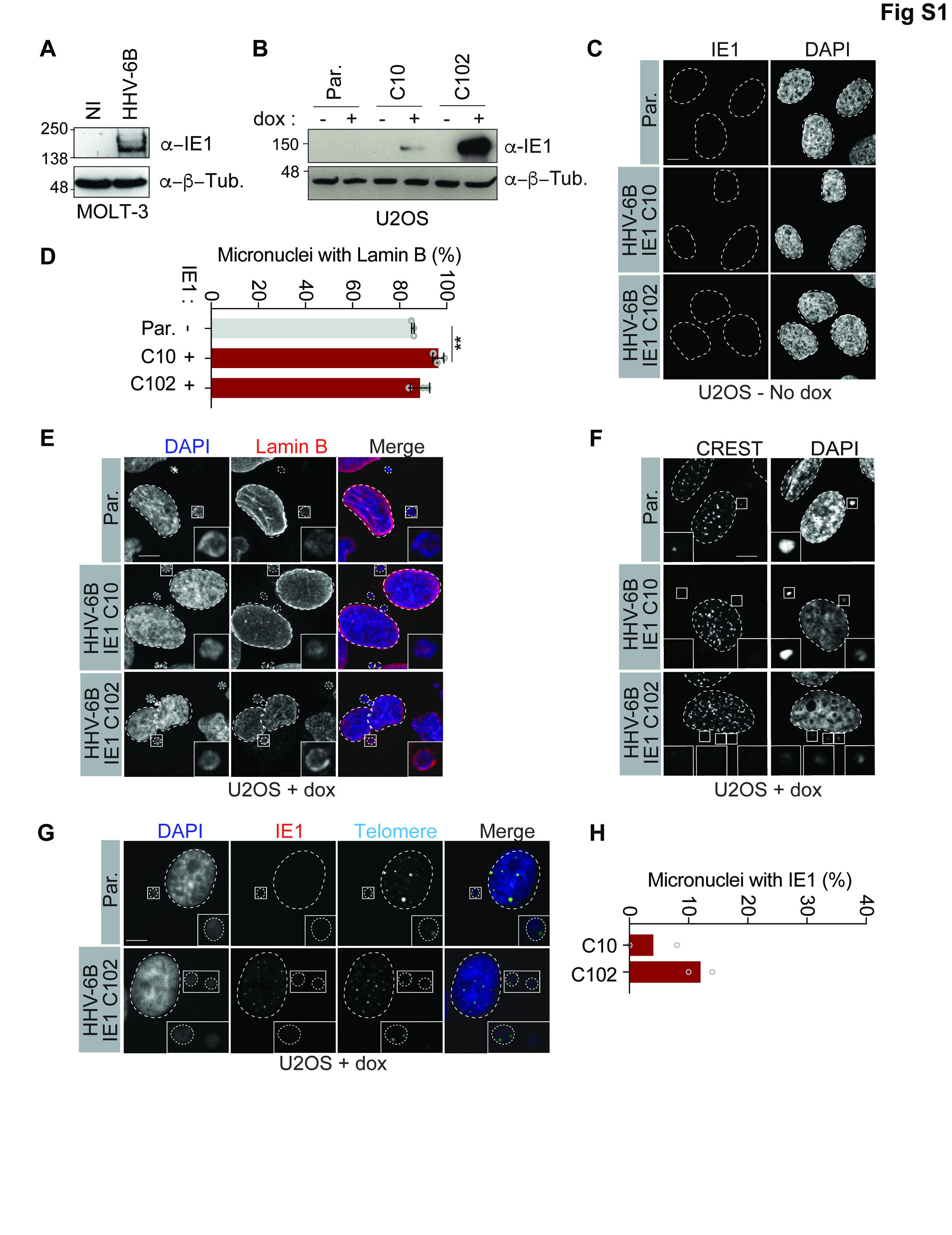

**Fig. S1. Characterization of MOLT-3 infected cells and U2OS clone C10 and C102 expressing IE1, related to Fig 1** *(A-B)* Whole cell extracts (WCEs) from infected MOLT-3 (*A*) and U2OS cells treated with or without 1 µg/mL Dox (*B*) were immunoblotted for IE1. β-tubulin (Tub.) was used as a loading control. NI: non-infected, Par.: parental cell line. *(C, F, G)* Representative images of parental (Par.) U2OS cells and IE1-expressing U2OS stable cell lines corresponding to Fig. 1*B*, *D,* and *E,* respectively. Cells were treated as described in Fig. 1*B* and either processed for IE1 (*C* (without DOX), *G*) or centromeres (CREST) immunofluorescence (*F*) or FISH for telomere detection (*G*). *(D-E)* Dox-inducible U2OS (Par.) and U2OS IE1 stable cell lines were treated as in Fig. *1B* and immunostained for lamin B. Quantification of lamin B-positive micronuclei is presented in (*D*) and representative images are in (*E*). In (*D*), data represent the mean ± SD (*n* = 3), ***p* ≤ 0.01. *(H)* Quantification of micronuclei colocalizing with IE1 foci. IE1-expressing cells were treated as in Fig. 1*C*, fixed, and immunostained for IE1. Data represent the mean ± SD (*n* = 2, > 100 nuclei/condition). Scale bar = 5 μm.

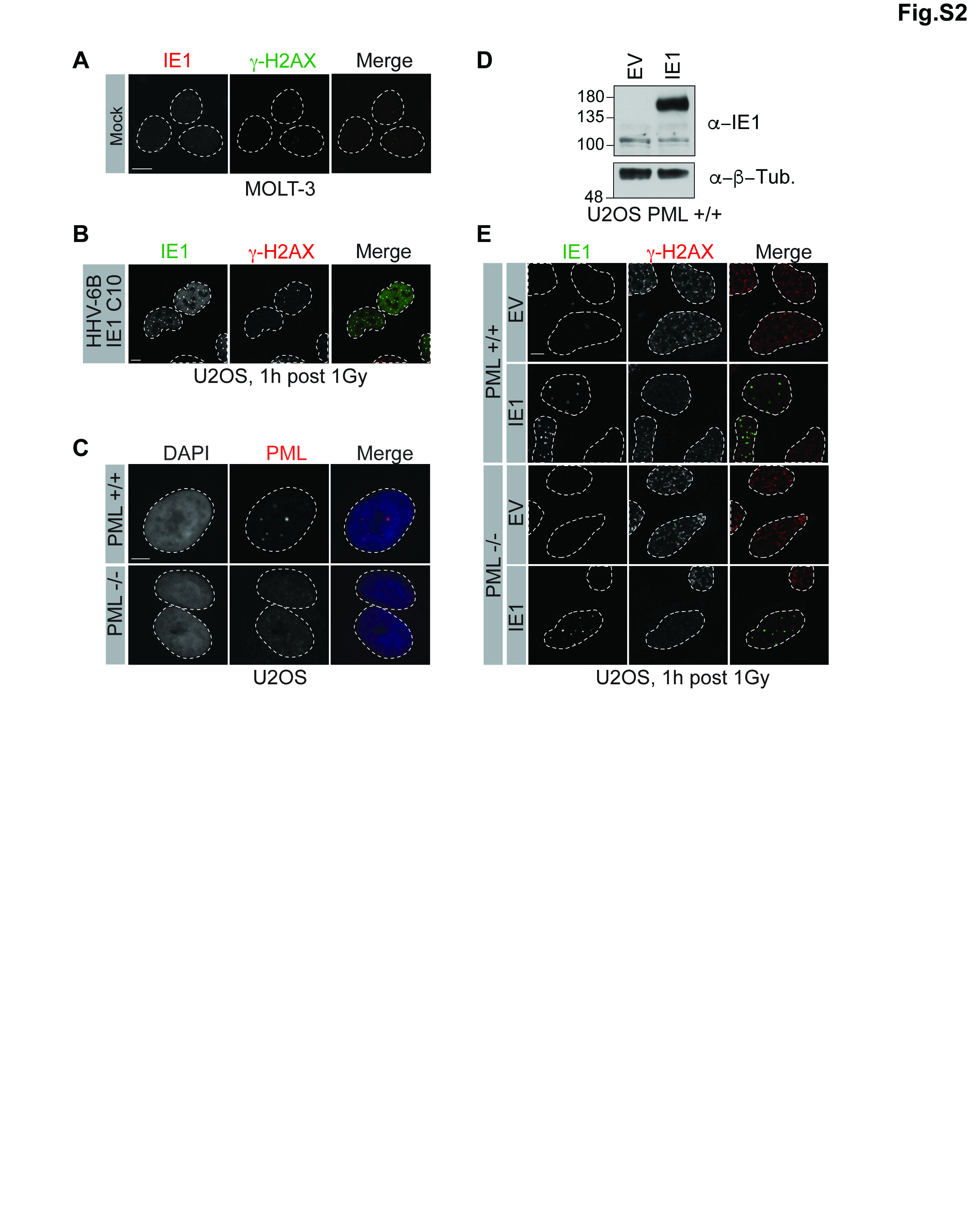

**Fig. S2. Representative images of the phenotypes quantified in MOLT-3 and U2OS cells and validation of transient expression of IE1 in U2OS cells, related to Fig 2** *(A)* Representative images of untreated MOLT-3 cells immunostained for γ-H2AX. Cells were treated as in Fig 2*A*. *(B)* Representative images of γ-H2AX in irradiated IE1-expressing U2OS cells (Clone C102). Cells were treated as described in Fig. 2*C*. *(C)* Representative images of untreated U2OS *PML*^+/+^ and ^-/-^ cells immunostained for PML. *(D)* Western blot analysis of U2OS cells transfected with untagged IE1 or an empty vector (EV). β-tubulin (Tub.) was used as a loading control. *(E)* Representative images of U2OS *PML*^+/+^ and ^-/-^ cells transiently transfected with untagged IE1 or an empty vector (EV). Cells were irradiated with 1 Gy, fixed 1 h post irradiation, and immunostained for IE1 and γ-H2AX (scale bars, 5 μm).

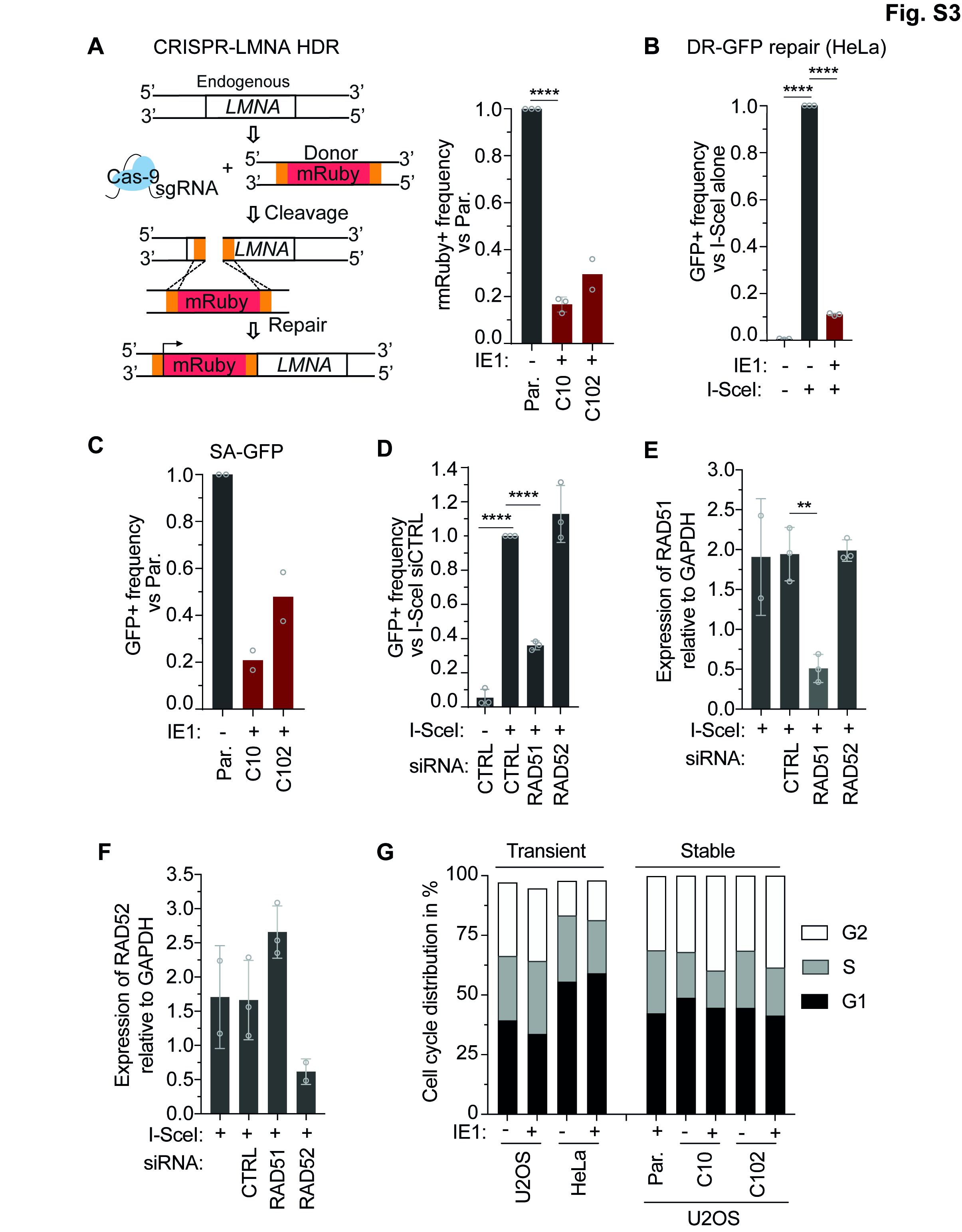

**Fig. S3.** **Characterization of additional DNA repair** **reporter assays and associated controls in cells expressing IE1, related to Fig. 3** *(A)* CRISPR-LMNA HDR assay in Dox-inducible IE1 U2OS SA-GFP stable cell lines. Cell lines were plated and induced for IE1 expression for 24 h prior to transfection with plasmids encoding Cas-9, a *LMNA* sgRNA, and the mRuby2-LMNA donor. The percentage of mRuby-positive cells was analyzed by flow cytometry 48 h post-transfection and normalized to the percentage of mRuby-positive U2OS SA cells (Par.) in each replicate. *(B)* DNA repair assays for homologous recombination in HeLa cells were performed as described for U2OS cells in Fig. 3*A*. *(C)* SA-GFP assays were analyzed in Dox-inducible IE1 U2OS SA-GFP stable cell lines. Cell lines were plated and induced for IE1 expression for 24 h prior to transfection with a plasmid encoding I-Sce1 endonuclease. The percentage of GFP^+^ cells was analyzed by flow cytometry 48 h post-transfection and normalized to the percentage of GFP-positive U2OS SA cells (Par.) in each replicate. Data represent the mean ± SD (*n* = 2). *(D)* Validation of the BIR repair assay using siRNAs against RAD51 and RAD52. A non-targeting siRNA (CTRL) was used as a control. *(E-F)* RAD51 (*E*) and RAD52 (*F*) RT-qPCR was performed on U2OS BIR cells. Expression was normalized against GADPH. *(G)* Cell cycle analyses of cells expressing IE1. The indicated cell lines were plated and either transfected or induced for IE1 expression for 24 h prior to fixation, PI staining, and flow cytometry (*n* = 1). Unless stated otherwise, data represent the mean ± SD (at least *n* = 3). ***p* < 0.01, *****p* < 0.0001.

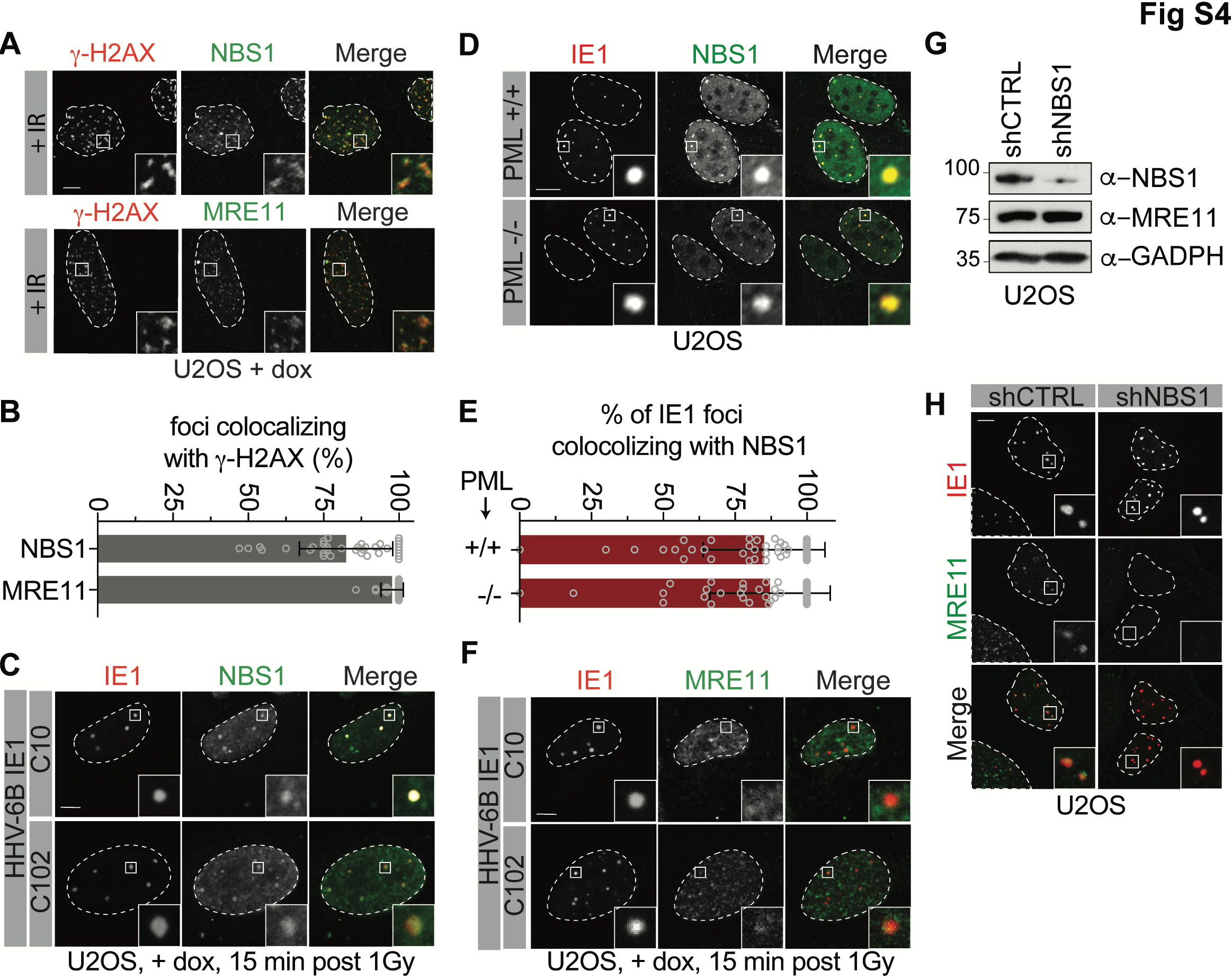

**Fig. S4**. **Representative images and analyses of the colocalization of IE1 with NBS1 in U2OS cells, related to Fig 4** *(A)* Representative images of the positive controls (irradiated cells) for Fig. 4*C* and *E*. Irradiated U2OS cells (+IR) were fixed 15 min post-irradiation (1 Gy) and immunostained for γ-H2AX, NBS1, or MRE11. *(B)* Quantification of γ-H2AX foci colocalized with NBS1 or MRE11 in irradiated cells, presented as a percentage of the total foci/cell colocalized with the indicated protein. Data represent the mean ± SD (*n* = 3). *(C, F)* Representative immunofluorescence of IE1-expressing U2OS stable cell lines induced with Dox and irradiated with 1 Gy. Cells were fixed 15 min post-irradiation and processed as described in Fig. 4*A*. (*D-E*) Representative immunofluorescence and quantification of IE1 foci that colocalize with NBS1 in U2OS *PML*^+/+^ and ^-/-^ cells transiently expressing untagged IE1. Data represent the mean ± SD (*n* = 3) *(G)* WCEs from U2OS shCTRL and shNBS1 stable cell lines were immunoblotted for NBS1 and MRE11. GAPDH was used as a loading control. *(H)* Representative images for the quantification presented in Fig. 4*F*. . Scale bars = 5 μm.

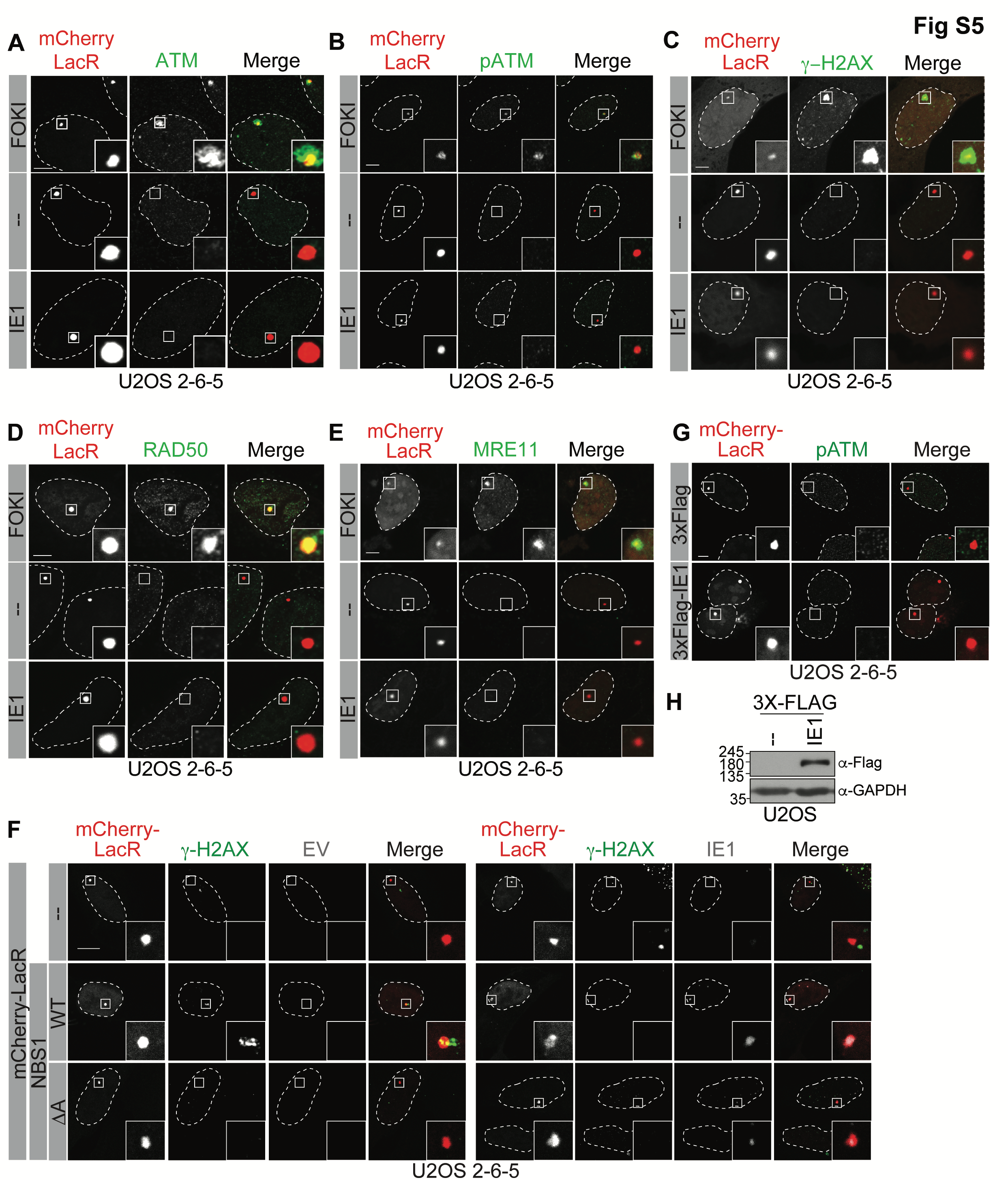

**Fig. S5. Representative image of the quantification presented in Fig. 5 and validation of Flag-IE1 expression in U2OS cells** *(A-E)* Representative images of the data presented in Fig. 5*B*. *(F)* Representative immunofluorescence used for the quantification presented in Fig. 5*D*. *(G)* Representative mCherry-LacR immunofluorescence used as a negative control in Fig. 5*E*. (*H*) WCEs from U2OS cells transiently transfected with the 3×FLAG-tagged IE1 expression vector or an empty vector were immunoblotted for FLAG. GAPDH was used as a loading control. Scale bars = 5 μm.

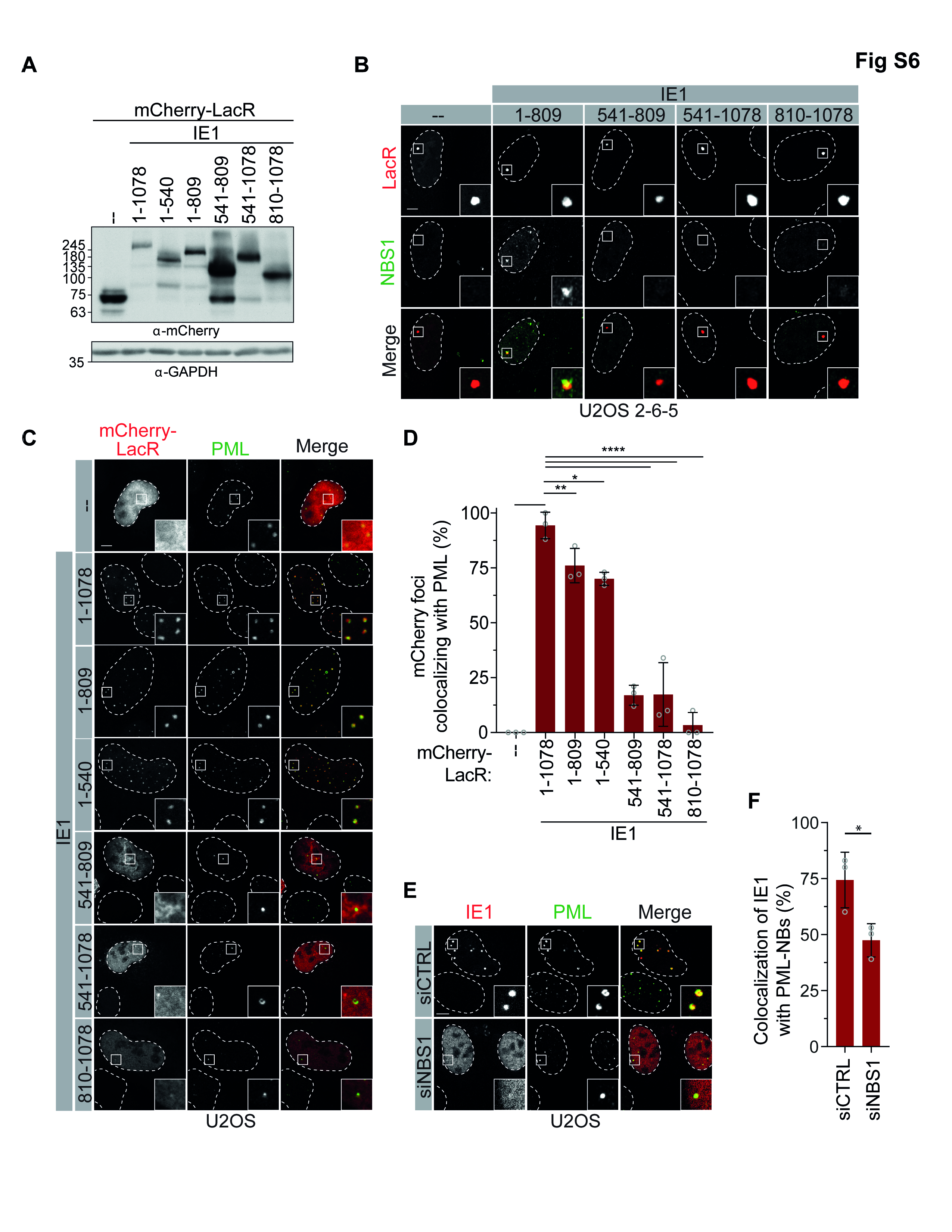

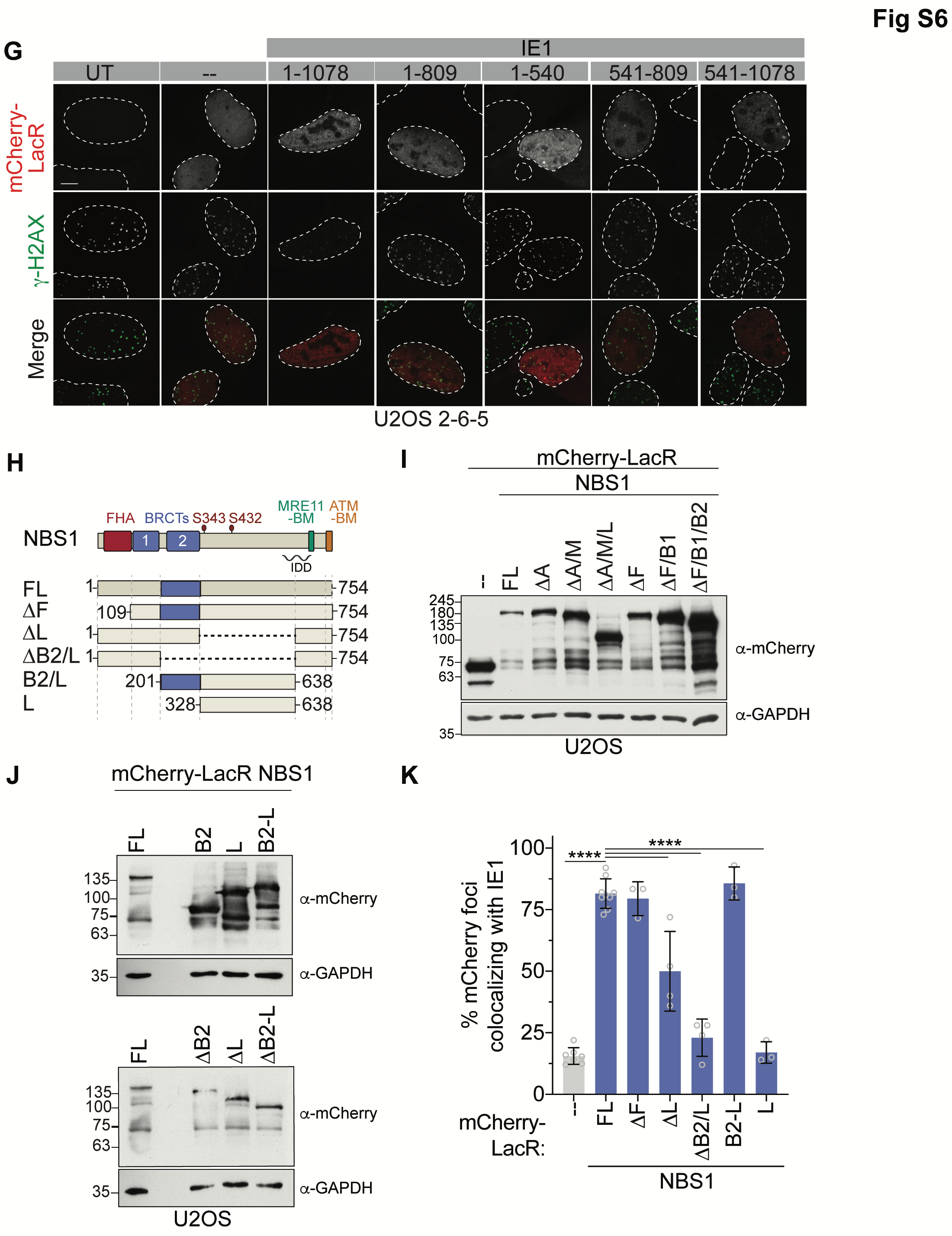

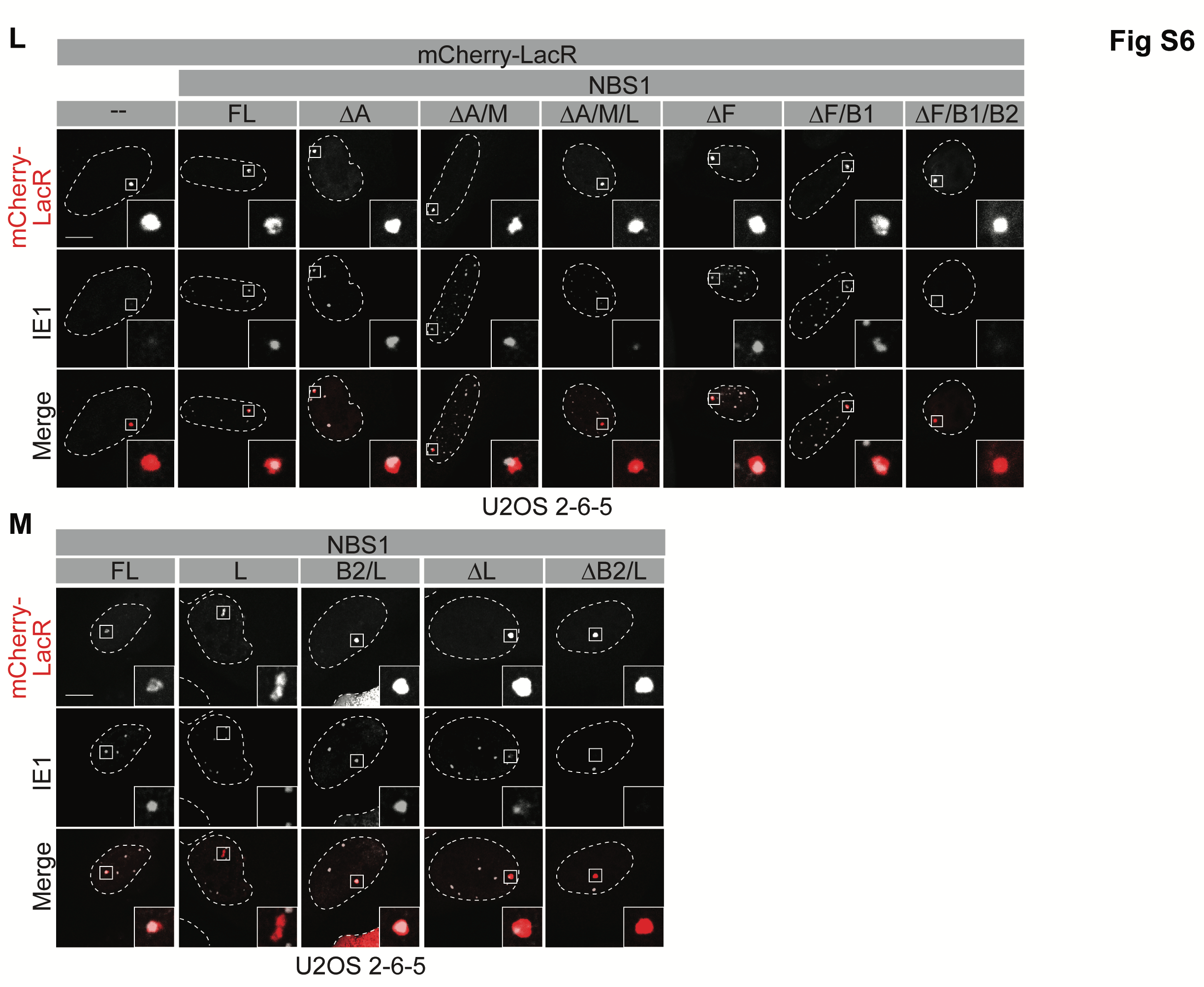

**Fig. S6. Validation of the expression and cellular localization of IE1 and NBS1 fragments used in this study, related to Fig6** *(A, I, J)* WCEs from U2OS cells transiently transfected with the indicated mCherry-LacR or 3×FLAG fusion proteins and treated with the indicated siRNAs were immunoblotted for mCherry or FLAG. GAPDH was used as a loading control. The mCherry-LacR backbone was used as a negative control (--) in (*A* and *I*). *(B, G, L, M)* Representative images used for the quantifications presented in Fig. 6*B (B)*, Fig. 6*D* *(G)*, Fig. 6*G,* and *SI Appendix,* Fig. S6*K* *(L, M)*. *(C-D)* Quantification and representative immunofluorescence of the minimal domain of mCherry-LacR-IE1 to be recruited to PML bodies. U2OS cells were transiently transfected with the indicated mCherry-LacR constructs and immunostained for PML. *(E-F)* Quantification and representative immunofluorescence of the colocalization of IE1 with PML nuclear bodies (NBs) in NBS1-depleted cells. U2OS cells were treated with the indicated siRNAs, transfected with an IE1-expressing vector and immunostained for PML. *(H)* Schematic of NBS1 and the protein fragments used in this study. FHA, forkhead-associated domain; BRCT, BRCA1 C-terminal domain; MRE11-BM, MRE11-binding motif; ATM-BM, ATM-binding motif; IDD, intrinsically disordered domain. *(K)* Quantification of the colocalization of the indicated mCherry-LacR-NBS1 constructs with IE1. Data represent the mean ± SD (*n* = 3). **p* < 0.05, ***p* < 0.01, *****p* < 0.0001. Scale bars = 5 μm.

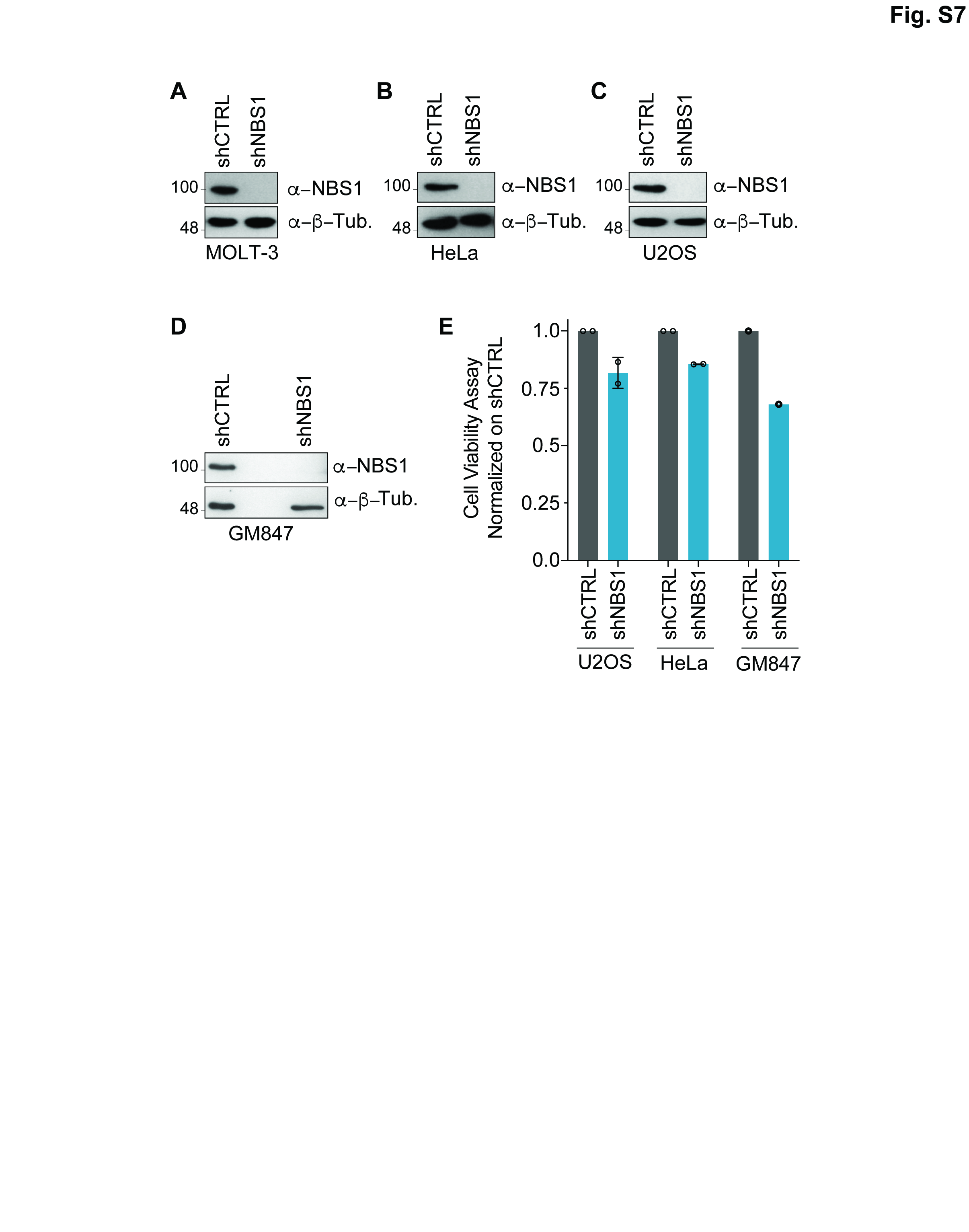

**Fig. S7. Validation of NBS1 knockdown and its impact on cell viability** *(A-D)* WCEs from MOLT-3 (*A*), HeLa (*B*), U2OS (*C*), and GM847 (*D*) cell lines expressing an shRNA against NBS1 (shNBS1) or a control shRNA (shCTRL) were immunoblotted for NBS1. β-Tubulin (Tub.) was used as a loading control. *(E)* Quantification of cell viability by CellTiter-Glo® assay. The numbers of viable cells were determined using standard curves for each cell line, with the number of cells in the shCTRL sample set to 1.0. Data represent the mean ± SD (*n* = 1 and *N* = 2).

**Table S1.** Plasmids used in this study

| **Plasmid** | **Source** |
| --- | --- |
| pcDNA4/TO/myc-His-HHV-6B IE1 | Previously described^2^ |
| pcDNA4/TO/myc-His | ThermoFisher, V103020 |
| pDEST-PB-TetO | Previously described^3^ |
| PB-CA-rtTA-IRES-NEO | Previously described^3^ |
| PB-TetO-HHV-6B IE1 | This study |
| pDEST-mCherry-LacRnls | Previously described^4^ |
| mCherry-LacRnls HHV-6B IE1 1-1078 | This study |
| mCherry-LacRnls HHV-6B IE1 1-809 | This study |
| mCherry-LacRnls HHV-6B IE1 1-540 | This study |
| mCherry-LacRnls HHV-6B IE1 541-809 | This study |
| mCherry-LacRnls HHV-6B IE1 541-1078 | This study |
| mCherry-LacRnls HHV-6B IE1 810-1078 | This study |
| pDEST-FRT-TO-3×FLAG | Gift from Anne-Claude-Gingras (V4978)^7^ |
| 3×FLAG HHV-6B IE1 1-1078 | This study |
| mCherry-LacRnls NBS1 1-754 (WT) | This study |
| mCherry-LacRnls NBS1 1-733 (∆A) | This study |
| mCherry-LacRnls NBS1 1-638 (∆A/∆M) | This study |
| mCherry-LacRnls NBS1 1-327 (∆A/∆M/∆L) | This study |
| mCherry-LacRnls NBS1 109-754 (∆F) | This study |
| mCherry-LacRnls NBS1 201-754 (∆F/∆B1) | This study |
| mCherry-LacRnls NBS1 328-754 (∆F/∆B1/∆B2) | This study |
| mCherry-LacRnls NBS1 ∆201-638 (∆B2/L) | This study |
| mCherry-LacRnls NBS1 ∆201-327 (∆B2) | This study |
| mCherry-LacRnls NBS1 ∆328-638 (∆L) | This study |
| mCherry-LacRnls NBS1 201-327 (B2) | This study |
| mCherry-LacRnls NBS1 328-638 (L) | This study |
| mCherry-LacRnls NBS1 201-638 (B2/L) | This study |
| pLXIN2-NBS1 | Gift from Cary A. Moody^6^ |
| pLKO.1-Scramble | Addgene 136035 |
| pLKO.1-shNBS1 | Gift from Cary A. Moody^6^ |
| pMD2.g | Addgene 12259 |
| psPAX | Addgene 12260 |
| pBIR-GFP | Addgene 49807 |
| pCBASceI | Addgene 26477 |
| pX330-LMNA-gRNA1 | Gift from Jean-Yves Masson^8^ |
| pCR2.a-mRuby2-LMNA-Donor | Gift from Jean-Yves Masson^8^ |
| pX330-7a sgRNA (EJ7-GFP reporter) | Addgene 113620 |
| pX330-7b sgRNA (EJ7-GFP reporter) | Addgene 113624 |

| **Table S2.** Primers, siRNA, shRNAs, and FISH probe sequences used in this study | | |
| --- | --- | --- |
| **Name** | **Sequence (5′-3′)** | **Purpose** |
| shCTRL  *(Sigma: SHC202)* | CCGGCAACAAGATGAAGAGCACCAACTCGAGTTGGTGCTCTTCATCTTGTTGTTTTT | Control shRNA in U2OS, MOLT-3, HeLa, GM847 cells |
| shNBS1  *(TRCN0000010393)* | GAAGAGTGGCTAAGGCAGGAA | KD NBS1 in U2OS, MOLT-3, HeLa, GM847 cells |
| U67-68 FW | TTCCGGTATATGACCTTCGTAAGC | qPCR and ddPCR |
| U67-68 RV | GATGTCTCACCTCCAAATCTTTAGAAAT | qPCR and ddPCR |
| U67-68 Probe | FAM-ACATTATAT-ZEN-RTCGAAYYTGACR CTACCTTCCG-IABkFQ  *Mixed probe where R = A or G, Y = C or T to accommodate the sequences of HHV-6A and -6B* | ddPCR^c^ |
| RPP30 FW | GATTTGGACCTGCGAGCG | qPCR and ddPCR |
| RPP30 RV | GCGGCTGTCTCCACAAGT | qPCR and ddPCR |
| RPP30 Probe | HEX-TCTGACCTG-ZEN-AAGGCTCTGCGCG-IABkFQ | ddPCR^c^ |
| GAPDH FW | CAACGTGTCAGTGGTGGACC | RT, qPCR^b^ |
| GAPDH RV | TCGTTGAGGGCAATGCCAGC | RT, qPCR^b^ |
| RAD51 FW | TCTCTGGCAGTGATGTCCTGGA | qPCR |
| RAD51 RV | TAAAGGGCGGTGGCACTGTCTA | qPCR |
| RAD52 FW | ATGCTTTGGACAGTGCCAGT | qPCR |
| RAD52 RV | CCAGCCATGCGGCTACTTAT | qPCR |
| siRAD52  *(custom)* | GGCUACAAUCUUCAUUUAAdTdT | KD in U2OS BIR assays |
| siRAD51  *(M-003530-04-0005)* | Smartpool:  D-003530-02: GAAGCUAUGUUCGCCAUUA  D-003530-05: GCAGUGAUGUCCUGGAUAA  D-003530-07: CCAACGAUGUGAAGAAAUU  D-003530-08: AAGCUAUGUUCGCCAUUAA | KD in U2OS BIR assays |
| siNBS1  (custom) | AACAUACGUAGCUGACACAGAdTdT | KD in U2OS immunoprecipitation/  immunostaining |
| siCTRL  *(D-001210-02-02)* | UAAGGCUAUGAAGAGAUAC | CTRL in U2OS BIR assays |
| Cy5-TelC PNA probe  *(F1003)* | [Cy5]-OO-CCCTAACCCTAACCCTAA | FISH |

^a^ KD: knockdown, gDNA: genomic DNA

^b^ Used to validate the efficiency of DNAse treatment prior to RT-PCR

^c^ FAM: fluorescein, HEX: HEX fluorophore, ZEN: ZEN^TM^ quencher, IABkFQ: 3′ Iowa Black® forward quencher

**Table S3.** Antibodies used in this study

| **Primary antibodies** | | |
| --- | --- | --- |
| **Target** | **Catalog number (Source)** | **Purpose^a^** |
| Mouse anti-phospho-Histone H2A.X (Ser139) (γ-H2AX), clone JBW30 | 05-636 (Millipore) | IF |
| Rabbit anti-phospho-Histone H2A.X (Ser139) (γ-H2AX) | 2577L (Cell Signaling Technology) | IF |
| Mouse anti-phospho-Histone H2A.X (Ser139) (γ -H2AX), clone JBW30, alexa fluor conjugate | 05-636-AF555 (Millipore) | IF |
| Mouse anti-β-tubulin | CP06 (Sigma) | IB |
| Mouse anti-GAPDH | 39-8600 (Invitrogen) | IB |
| Rabbit anti-IE1 | Homemade^1^ | IF, IB |
| Rabbit anti-IE1, alexa fluor conjugate | Homemade^1^ | IF |
| Rabbit anti-lamin B1 | Ab16048 (Abcam) | IF |
| Human CREST anti-centromere serum | HCT-0100 (Immunovision) | IF |
| Mouse anti-PML (PG-M3) | sc-966 (Santa Cruz Biotechnology) | IF |
| Mouse anti-NBS1 1D7 | GTX70224 (GeneTex) | IF, IB |
| Mouse anti-MRE11 12D7 | GTX70212 (GeneTex) | IF, IB |
| Mouse anti-RAD50 13D3 | GTX70228 (GeneTex) | IF, IB |
| Rabbit anti-FLAG | 2368S (Cell Signaling Technology) | IF, |
| Mouse anti-FLAG M2 | F1804 (Millipore Sigma) | IF, IB, IP |
| Rabbit anti-ATM | A300-299A (Bethyl Laboratories) | IF, IB |
| Rabbit anti-pATM (S1981) D25E5 | 13050 (Cell Signaling Technology) | IF, IB |
| Rabbit anti-mCherry | NBP2-25157 (Novus) | IB |
| **Secondary antibodies** | | |
| **Target** | **Catalog number (Source)** | **Purpose^a^** |
| Alexa Fluor 488 goat anti-rabbit | A-11034 (Thermo Fisher Scientific) | IF |
| Alexa Fluor 555 goat anti-rabbit | A-21428 (Thermo Fisher Scientific) | IF |
| Alexa Fluor 647 goat anti-rabbit | A-21244 (Thermo Fisher Scientific) | IF |
| Alexa Fluor 488 goat anti-mouse | A-11029 (Thermo Fisher Scientific) | IF |
| Alexa Fluor 555 goat anti-mouse | A-21424 (Thermo Fisher Scientific) | IF |
| Alexa Fluor 647 goat anti-mouse | A-21236 (Thermo Fisher Scientific) | IF |
| Alexa Fluor 647 donkey anti-human | A-21445 (Thermo Fisher Scientific) | IF |
| Goat anti-rabbit HRP | 7074 (Cell Signaling Technology) | IB |
| Sheep anti-mouse-HRP | A-9044 (Sigma) | IB |

^a^ IF: immunofluorescence, IB: immunoblotting, IP: immunoprecipitation
